# Small brown planthopper infestation enhances it reproduction and insecticide tolerance by manipulating glucose distribution and levels in rice

**DOI:** 10.64898/2026.01.23.701313

**Authors:** Hainan Zhang, Qi Zhang, Huichen Ge, Jiaping Wei, Kun Qian, Xiaolong Liu, Hai Li, Jianjun Wang

## Abstract

The evolutionary arms race between plants and insects involves not only direct defense and counter-defense but also sophisticated resource manipulation. However, how herbivorous insects respond to host nutritional signals to modulate their fitness traits remains unclear. This study investigates how the small brown planthopper (SBPH, *Laodelphax striatellus*) manipulates host plant carbohydrate allocation, and to elucidate the molecular mechanisms by which the acquired glucose enhances SBPH fecundity and insecticide tolerance. Using molecular, pharmacological, and biochemical approaches, we found that SBPH infestation induced systemic carbohydrate reallocation in rice, elevating whole-plant glucose levels by promoting aerial accumulation while depleting root reserves. Host-derived glucose enhanced SBPH fecundity by activating the target of rapamycin (TOR) pathway, upregulating juvenile hormone (JH) signaling, and increasing vitellogenin production. For imidacloprid tolerance, glucose boosted glutathione S-transferase (GST) activity via two synergistic mechanisms: by upregulating glutamate cysteine ligase (GCL) to increase glutathione synthesis, and transcriptionally via the glucose-TOR-JH axis to induce *LsGSTe1* (SBPH epsilon class GST) and *LsGSTo1* (SBPH omega class GST) expression. Our findings establish host-derived glucose as a central signaling molecule that SBPH utilizes to modulate conserved pathways for simultaneous optimization of reproduction and insecticide resistance. This reveals a multifaceted nutrient-responsive mechanism in insect pests and identifies the glucose-TOR-JH axis as critical molecular targets for developing nutrient-based pest control strategies, such as disrupting insect nutrient-sensing pathways or modulating host carbohydrate metabolism.

**Highlights:**

1. SBPH infestation elevates whole-plant glucose levels by promoting aerial accumulation while suppressing root abundance.
2. SBPH-induced glucose in the rice aerial tissues boosts reproduction of SBPH via TOR-JH-Vg pathway.
3. SBPH-induced glucose in the rice aerial tissues enhances imidacloprid tolerance of SBPH through GCL-GSH-GST and TOR-JH-GST axis.

**Graphical Abstract:** 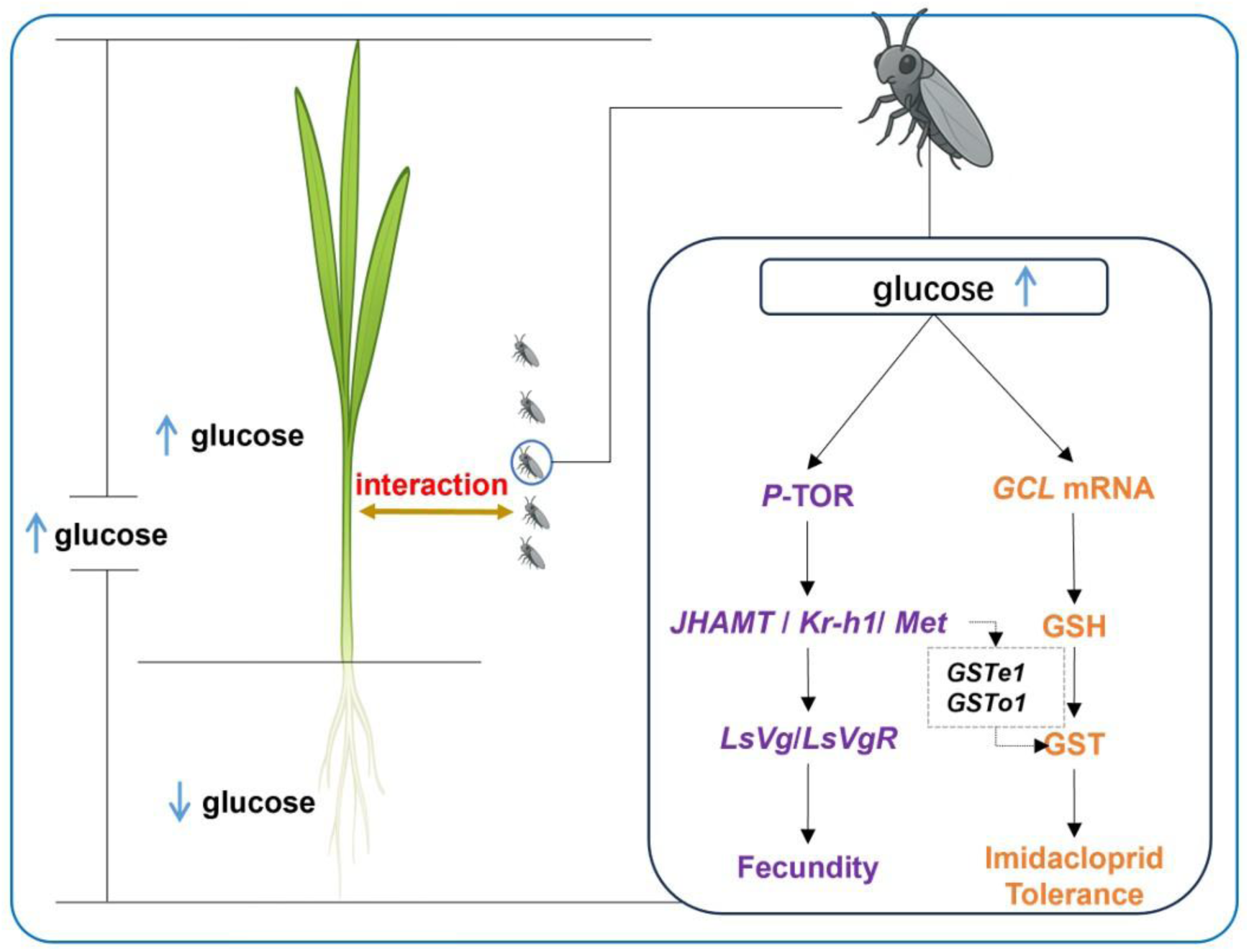

## Introduction

The co-evolutionary arms race between plants and insect herbivores involves complex interactions where defense mechanisms are met with sophisticated counter-strategies. While plants employ diverse defense mechanisms, including chemical deterrents and physical barriers, to resist insect herbivores, insects counteract these defenses through effector proteins and detoxifying enzymes^1–5^. Beyond these direct confrontations, emerging evidence highlights a more subtle battleground in the form of resource manipulation, wherein both plants and insects manipulate carbohydrate allocation to gain ecological advantages, and insect herbivores depend on host-derived carbohydrates to complete their life cycles^6–9^. Nevertheless, the key metabolic currencies targeted by herbivores, and how these plant-supplied metabolites drive insect physiological adaptation, remain unclear. Among plant primary metabolites, glucose occupies a unique position as both the transportable carbon source and a conserved signaling molecule across kingdoms^10–12^. In plants, glucose signaling modulates development, stress responses, and hormone signaling^13,14^, whereas in insects, it modulates diverse physiological processes such as chitin biosynthesis^15^, metamorphosis^10,16^, reproduction^17–19^, and stress tolerance^20^. Despite this centrality, the mechanisms through which glucose mediates plant–insect interactions, particularly how insects utilize host glucose to enhance fitness, remain elusive.

Fecundity is a key determinant of insect population dynamics, and juvenile hormone (JH) is well established as a master regulator of insect reproduction^21^. The target of rapamycin (TOR) pathway, a conserved serine/threonine kinase that integrates nutrient availability with developmental and metabolic homeostasis^22^, has been shown to link nutrient signals to JH biosynthesis and vitellogenin (Vg) production in several insect species^23–25^. Notably, TOR activation by glucose is evolutionarily conserved in mammals and plants^22,26–28^, and sugar-promoted TOR activation has also been reported in *Drosophila*²⁹, raising the hypothesis that glucose may regulate insect fecundity via the TOR-JH-Vg axis. However, direct evidence for glucose-mediated TOR activation in hemipteran insects and its functional connection to JH signaling and reproduction is lacking, representing a critical gap in our understanding of nutrient-driven insect adaptation.

In addition to fecundity, insecticide tolerance is another major factor shaping insect population dynamics and pest management outcomes. While the role of glucose in insecticide tolerance remains unclear, accumulating evidence suggests that carbohydrates can enhance stress resistance in insects, as evidenced by the reports that a high-sucrose diet upregulates *glutathione S-transferase* (*GST*) genes and confers malathion resistance in *Bactrocera dorsalis*^30^, and sucrose-fed *Drosophila melanogaster* exhibits elevated GST activity and enhanced tolerance to multiple stresses^31^. Since sucrose is hydrolyzed into glucose in insects^32^, these findings imply a potential role for glucose in mediating insecticide detoxification. Mechanistically, glucose has been shown to upregulate glutamate cysteine ligase (GCL), the rate-limiting enzyme in glutathione (GSH) synthesis^33,34^, via insulin signaling in mammals^35^, and GSH is an essential co-substrate for GST-mediated detoxification^36^. Furthermore, both TOR and JH have been implicated in regulating GST activity^37–39^, suggesting that glucose may modulate insecticide tolerance through dual GCL-GSH and TOR-JH pathways. However, the existence and functional relevance of these glucose-dependent regulatory networks in insect detoxification remain unvalidated.

Rice (*Oryza sativa*), a global staple crop critical for food security^40^, faces severe yield losses due to infestations by insect pests including the small brown planthopper (SBPH, *Laodelphax striatellus*)^41^. SBPH populations have developed some degree of insecticide resistance while maintaining high reproductive rates, leading to severe infestations^41^. However, it remains unclear how glucose metabolism in rice plants responds to SBPH infestation, and how such metabolic changes influence key SBPH fitness parameters, particularly reproduction and pesticide tolerance. In this study, we found that infestation of SBPH orchestrates a systemic reallocation of host carbohydrates, culminating in glucose-enriched aerial tissues. Notably, SBPH capitalizes on this nutritional change to fuel reproduction through the glucose-TOR-JH-Vg axis and to fortify detoxification by upregulating GST activity via dual metabolic (GCL-GSH) and regulatory (TOR-JH) pathways. These findings not only provide direct evidence of the involvement of glucose-responsive TOR signaling in plant-insect interactions, but also establish a molecular foundation for developing nutrient-based pest control strategies.

## Results

### SBPH infestation systemically alters glucose distribution and levels in rice

To assess the impact of SBPH feeding on rice carbohydrate metabolism, we first monitored glucose dynamics in aerial tissues over time. The results showed that after infestation with 25 third-instar nymphs per plant, glucose levels increased significantly by 27.70%, 72.43%, and 69.77% at 1, 3, and 5 days post-infestation, respectively (Figure 1A). Furthermore, this induction was also density-dependent, while a single nymph caused no detectable change, infestations with five or more nymphs for 5 days significantly elevated aerial glucose levels by 1.29- to 1.71-fold (Figure 1B). Specifically, the response peaked after infestations with 10–25 nymphs, with glucose concentrations plateauing between 5.10 – 5.27 μmol/g (Figure 1B). However, at the highest density of 30 nymphs, glucose levels declined to 4.86 μmol/g (Figure 1B), suggesting a potential stress threshold or altered plant response under extreme pest pressure. We next quantitatively compared the impact of different SBPH life stages. Infestation by virgin females, males, or nymphs (25 insects/plant) resulted in comparable glucose elevations (1.70-, 1.68-, and 1.65-fold, respectively). Strikingly, gravid females provoked a distinctly stronger response, with a 2.14-fold increase (Figure 1C). These results support a two-layer mechanism whereby a baseline glucose accumulation from probing or salivary effectors common to all life stages is then potentiated by gravid female-specific factors, most likely oviposition behavior, leading to more profound metabolic reprogramming.

**Figure 1.**
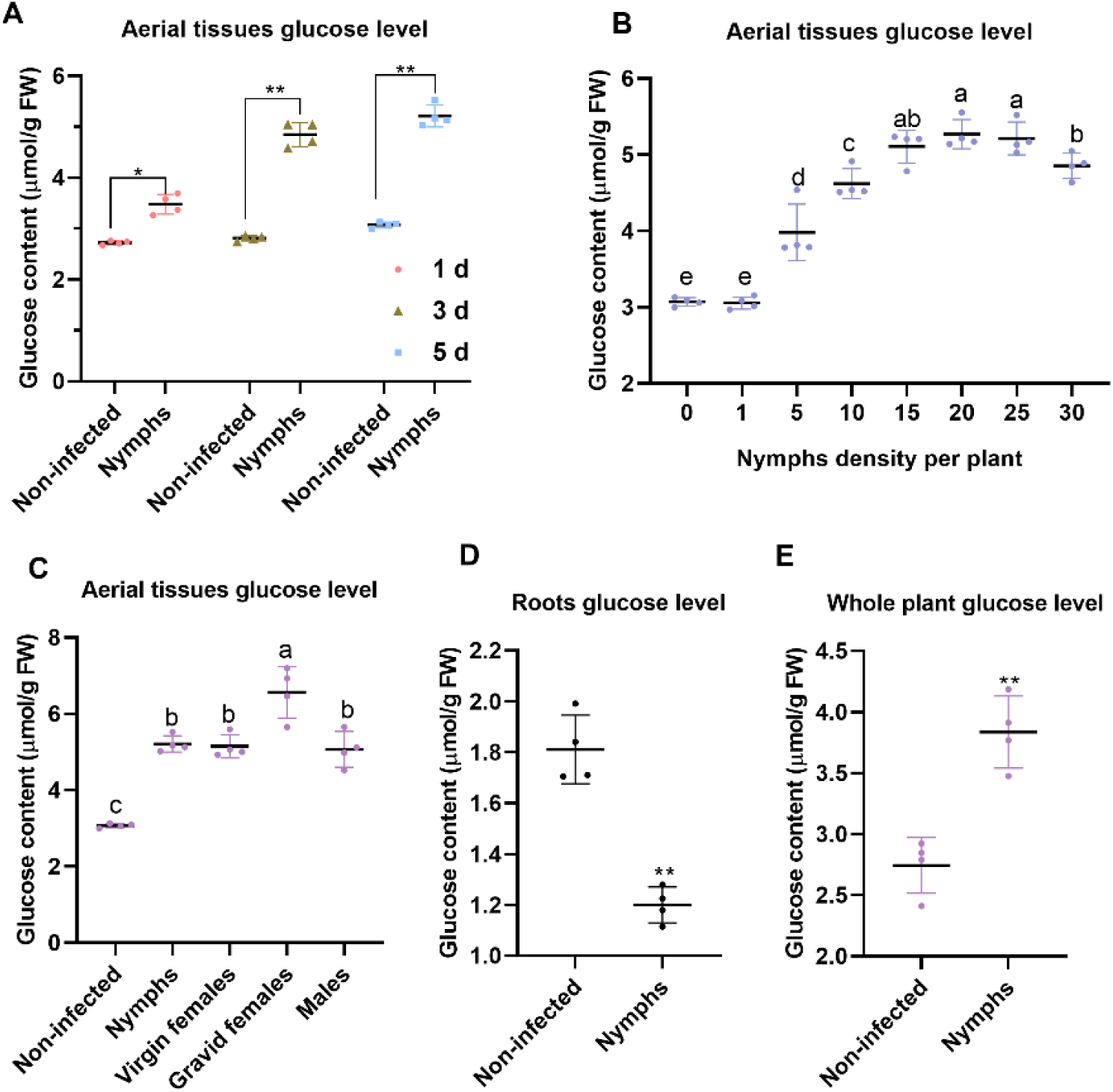
The effects of SBPH infestation on glucose distribution and levels in rice plants. (A, B, C) Glucose levels in aerial tissues of rice plants following SBPH infestation. (A) Rice plants infested with 25 third-instar nymphs/plant for 1, 3, 5 days. (B) Rice plants infested with third-instar nymphs at different density (1–30 insects/plant) for 5 days. (C) Rice plants infested with different developmental stages/sexes (third-instar nymphs, virgin females, gravid females, or males; 25 insects/plant) for 5 days. Uninfested plants served as controls. (D) Glucose levels in root tissues of the rice plants infested with 25 third-instar nymphs/plant for 5 days. (E) Whole-plant glucose levels of the rice plants infested with 25 third-instar nymphs/plant for 5 days. Data are from 4 independent biological replicates and presented as mean ± standard deviation (s.d). (B, C) Significant differences (one-way ANOVA with Tukey’s test; *p* < 0.05) are indicated by lowercase letters. (A, D, E) Were analyzed by Student’s *t*-test (**\****p* < 0.05, **\*\****p* < 0.01, ns = not significant). FW, fresh weight.

To determine whether the aerial glucose increase reflected a localized or systemic shift in carbohydrate allocation, we measured glucose levels in roots and whole plants after a 5-day infestation by 25 third-instar nymphs. In contrast to shoot accumulation, root glucose levels decreased by 33.78% (1.20 vs. 1.81 μmol/g; Figure 1D). Despite this reduction, whole-plant glucose content increased by 1.40-fold (Figure 1E). Together, these findings show that SBPH infestation systemically perturbs rice carbohydrate metabolism, enhancing overall glucose levels while redirecting its distribution to favor aerial tissues at the expense of root reserves.

### SBPH-induced elevation of rice glucose enhances fecundity and imidacloprid tolerance

To determine whether SBPH-induced metabolic changes in rice influence insect fitness, we employed a pre-infestation assay. When third-instar-nymphs were reared for 5 days on plants pre-infested (25 third-instar nymphs/plant for 3 days), their whole-body glucose levels increased by 20.08%, rising further to 27.32% upon emergence as female adults (Figure 2A). This was accompanied by a 25.39% increase in fecundity (from 97.17 to 121.83 eggs/female; Figure 2B) and significant upregulation of the vitellogenin gene *LsVg* and its receptor *LsVgR* (1.99- and 1.53-fold, respectively; Figure 2C), indicating that pre-infestation enhances reproductive output. Direct injection of glucose into nymphs recapitulated this phenotype, increasing fecundity by 31.53% (from 104.67 to 137.67 eggs/female; Figure 2D) and upregulating *LsVg/LsVgR* expression by 1.95- and 2.30-fold, respectively (Figure 2E), confirming that elevated glucose underlies the enhanced fecundity.

**Figure 2.**
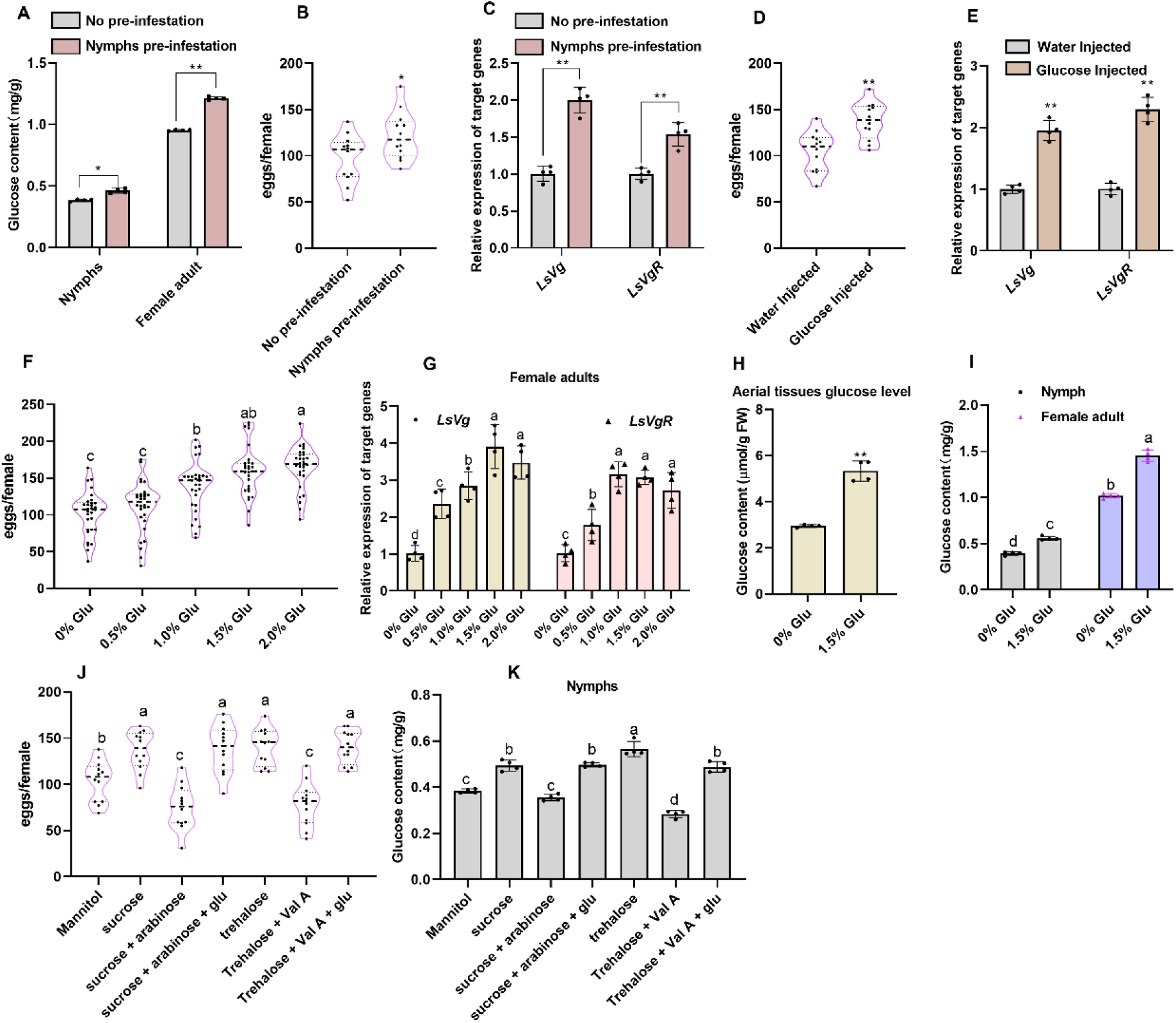
Aerial tissues glucose levels induced by SBPH infestation affect the fecundity. (A, B, C) Glucose levels, fecundity, and transcriptional responses of *LsVg* and *LsVgR* in SBPH on pre-infested rice plants. Third-instar nymphs were reared on rice plants that had been pre-infested with 25 nymphs for 3 days; control insects were fed on non-infested plants. (A) Glucose levels in nymphs (at 5 days post-infestation) and in females (25 insects/group). (B) Oviposition rate (1♀:1♂ per group). (C) Expression levels of *LsVg* and *LsVgR* in females (25 insects/group). (D) Oviposition rate (1♀:1♂ per group) after exogenous glucose injection into nymphs. (E) Expression of *LsVg* and *LsVgR* in female that developed from nymphs injected with exogenous glucose (25 insects/group). (F, G) Effects of glucose level on oviposition rate and gene expression in SBPH. Third-instar nymphs were reared on rice plants cultivated in IRRI solution supplemented with 0–2.0% glucose. (F) Oviposition rate (1♀:1♂ per group). (G) *LsVg* and *LsVgR* expression in female (25 insects/group). (H) Glucose content in aerial parts of rice plants cultured for 5 days in IRRI solution with 0% (control) or 1.5% glucose (1 plant/group). (I) Glucose levels in nymphs (5 days post-infestation) and adult females reared on 1.5% glucose-supplemented rice; controls fed on plants cultured in standard IRRI solution (25 insects/group). (J) Effect of dietary sugars on SBPH fecundity. Rice roots were irrigated with an IRRI solution containing mannitol, sucrose, or trehalose, each isotonic with 1.5% glucose, and nymphs were reared on these plants. Parallel treatment groups include nymphs from the sucrose- or trehalose-reared groups were injected with the sucrase inhibitor arabinose or the trehalase inhibitor Validamycin A (Val A), respectively, with or without a subsequent glucose injection. After eclosion, adults were paired (1♀:1♂ per group) and continued to be reared on correspondingly treated plants. Fecundity was measured as the total number of eggs laid per female. (K) Glucose levels in nymphs after 5 days of feeding under the treatments described in (J) (25 insects/group). Panel images (A, C, E, G, H, I, K) represent data from four biological replicates, while (B, D, J) and (F) include twelve and thirty replicates, respectively. All data are presented as mean ± s.d. (A-E, H) Were analyzed by Student’s *t*-test (**\****p* < 0.05, **\*\****p* < 0.01, ns = not significant). (F, G, I-K) Significant differences (one-way ANOVA with Tukey’s test; *p* < 0.05) are indicated by lowercase letters. Glu, Glucose.

To further validate this under more natural conditions, rice plants were cultivated in IRRI nutrient solution supplemented with glucose. Concentrations of 1.0–2.0% glucose significantly increased SBPH fecundity (138.8–164.8 eggs/female) and robustly upregulated *LsVg/LsVgR* by 2.34- to 3.91-fold and 1.78- to 3.15-fold, respectively, whereas 0.5% glucose was ineffective (Figure 2F, G). The 1.5% glucose treatment was selected for subsequent experiments, as it elevated fecundity to a level statistically equivalent to the 2.0% group. We confirmed that root-applied 1.5% glucose was translocated to aerial tissues, increasing glucose levels by 1.81-fold (Figure 2H) and subsequently increasing glucose content in feeding nymphs and adults by 42.01% and 42.77%, respectively (Figure 2I). These results establish that root glucose supplementation effectively mimics infestation-induced host alterations and enhances SBPH reproduction.

To exclude potential confounding effects of osmolarity, a mannitol solution equimolar to 1.5% glucose (83.3 mM) was used as an osmotic control. Since SBPH fecundity on mannitol-treated seedlings did not differ significantly from the 0% glucose control (Figure 2F, 2J), we concluded that the enhanced fecundity was not attributable to osmotic effects. In contrast, root treatment with an IRRI solution containing 83.3 mM sucrose or trehalose significantly enhanced fecundity relative to the mannitol control (103.67 eggs/female), with increases of 31.75% (136.58 eggs/female) and 35.28% (140.25 eggs/female), respectively (Figure 2J). Concurrently, these treatments also raised nymphal glucose content by 28.62% (0.494 mg/g) and 47.28% (0.565 mg/g), respectively, compared to mannitol control (0.384 mg/g) (Figure 2K). Since both disaccharides are hydrolyzed into glucose in insects, we hypothesized that their effect is glucose-dependent. Confirming this, inhibition of sucrase with arabinose^42^ or inhibition of trehalase with Validamycin A^43^ in the respective treatment groups reduced nymphal glucose content to 0.355 mg/g and 0.283 mg/g (Figure 2K), and decreased fecundity by 27.81% (74.83 eggs/female) and 24.76% (78.00 eggs/female) relative to mannitol (Figure 2J). A subsequent rescue experiment via glucose injection successfully restored both glucose levels and fecundity (Figure 2J, 2K). Collectively, these results demonstrate that SBPH enhance their fecundity by modulating rice glucose metabolism.

We then investigated whether host glucose also influences insecticide tolerance. Bioassays showed that the LC₅₀ of imidacloprid against third-instar nymphs (5.85 mg/L) after 5 days of exposure was significantly higher on rice plants pre-infested (25 third-instar nymphs/plant for 3 days) compared to those on non-pre-infested plants (2.26 mg/L; Table S1). Mirroring this, glucose supplementation (0.5–2.0%) induced a dose-dependent increase in LC₅₀ (2.35 to 6.46 mg/L; Table S1), indicating that elevated host glucose levels enhance insecticide tolerance. While the mannitol had no significant effect (LC₅₀ = 1.93 mg/L), ruling out the influence of rice plants under this osmotic pressure on SBPH’s tolerance to imidacloprid (Table S2), treatments with sucrose or trehalose markedly increased tolerance (LC₅₀ = 6.17 and 7.00 mg/L, respectively; Table S2), but this effect was abolished by inhibiting their respective hydrolases (LC₅₀ = 2.30 and 1.60 mg/L; Table S2). Critically, glucose supplementation via injection fully restored tolerance in arabinose- or Validamycin A-treated insects (Table S2), indicating that the observed tolerance, like fecundity, is contingent on glucose availability. In summary, SBPH infestation reprograms host glucose metabolism, and this nutritional change is utilized by the insect to simultaneously amplify its reproductive capacity and bolster its tolerance to imidacloprid.

### SBPH-induced host glucose enhances its fecundity via the TOR-JH-Vg axis

Given the critical role of JH in SBPH reproduction, we examined its involvement in the glucose-mediated fecundity increase. Pre-infestation significantly elevated JH III titers in both nymphs (33.03%, from 3.96 to 5.27 pg/insect) and newly emerged female adults (26.49%, from 6.11 to 7.72 pg/insect) (Figure 3A). Consistent with this, the expression levels of *JHAMT* (*juvenile hormone acid methyltransferase)*, as well as JH response genes *Kr-h1* (*Krüppel homolog 1*) and *Met* (*methoprene-tolerant*) were upregulated by 2.00-, 1.72-, and 1.46-fold, respectively, whereas transcripts of JH degradation enzymes (*JHE*, *juvenile hormone esterase*; and *JHEH*, *juvenile hormone epoxide hydrolase*) remained unchanged (Figure 3B), indicating a specific promotion of JH synthesis. This effect was recapitulated by supplementing rice with 1.5% glucose (Figure 3C), directly linking host glucose to JH pathway. To establish a functional relationship, we performed RNAi against *JHAMT*, *Kr-h1*, or *Met*. Silencing these genes (53.97–64.96% efficiency; Figure 3D) markedly reduced *LsVg* (61.92–72.28%) and *LsVgR* (62.17–75.82%) expression (Figure 3F), leading to a 51.21–54.02% decrease in egg-laying (50.6–53.7 vs. 110.1 eggs/female in dsEGFP controls; Figure 3E), confirming that glucose-enhanced reproduction depends on JH biosynthesis and signaling.

**Figure 3.**
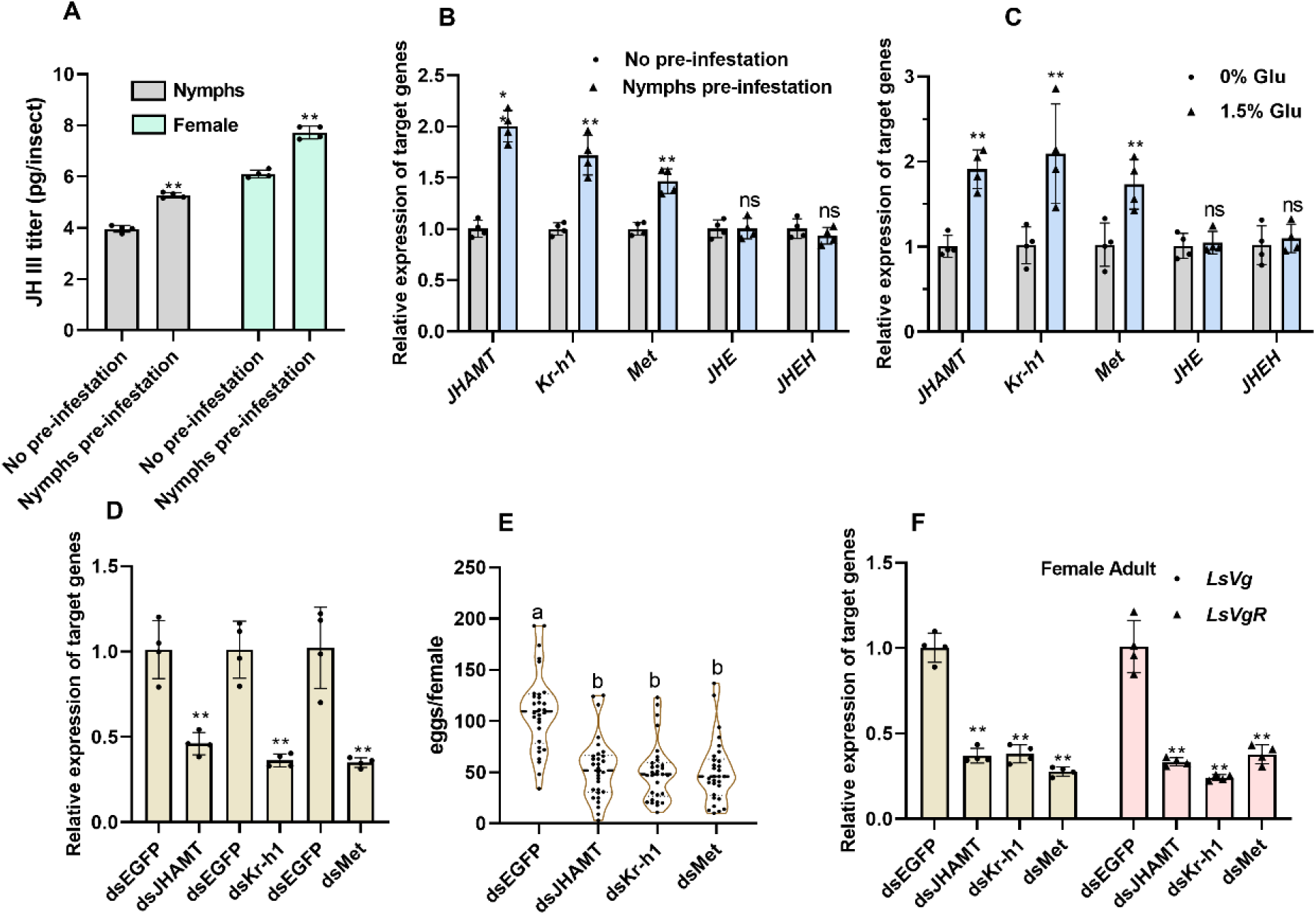
Glucose levels induced by SBPH infestation influence fecundity via the JH pathway. (A) Third-instar nymphs were fed for 5 days on rice plants that had been pre-infested with 25 nymphs for 3 days; JH III levels were measured in these nymphs and in the resulting adult females after eclosion. Control insects were fed on non-infested plants (25 insects/group). (B) Expression of JH-related genes (*JHAMT*, *Kr-h1*, *Met*, *JHE*, *JHEH*) in nymphs from the treatment described in (A) after 5 days of feeding. (C) Relative mRNA expression of JH pathway genes (*JHAMT*, *Kr-h1*, *Met*, *JHE*, *JHEH*) in nymphs after 5-day feeding on rice plants cultured in 1.5% glucose-supplemented IRRI solution. Control plants were grown in standard IRRI solution (25 insects/group). (D) Relative expression of *JHAMT*, *Kr-h1*, and *Met* of the third-instar nymphs were injected with corresponding dsRNAs and reared on rice cultured in standard IRRI solution for 5 days; dsEGFP served as control (25 insects/group). (E) Oviposition rate of adults eclosing from RNAi-treated nymphs in (D) (1♀:1♂ per group). (F) Expression of *LsVg* and *LsVgR* in females from RNAi-treated nymphs in (D) (25 insects/group). Panel images (A, B, C, D, F) represent data from four biological replicates, while (E) include thirty replicates. All data are presented as mean ± s.d. (A, B, C, D, F) Were analyzed by Student’s *t*-test (**\****p* < 0.05, **\*\****p* < 0.01, ns = not significant). (E) Significant differences (one-way ANOVA with Tukey’s test; *p* < 0.05) are indicated by lowercase letters.

We next sought to identify the upstream regulator linking glucose to JH signaling. Although total *TOR* mRNA levels remained unchanged (Figure 4A, 4C), supplementing rice with 1.5% glucose significantly enhanced TOR phosphorylation at Ser2448 in both nymphs and female adults (Figure 4B, 4D), indicating that glucose mediates activation of TOR at the post-translational level. Subsequent experiments showed that *TOR* knockdown (74.54% efficiency; Figure 4E) suppressed *LsVg* and *LsVgR* expression by 72.78% and 76.04%, respectively (Figure 4F), and reduced fecundity by 51.76% (53.58 vs. 111.08 eggs/female; Figure 4I), establishing critical role of TOR in glucose-mediated fecundity. Crucially, silencing *TOR* not only reduced JH III titers by 38.05% (from 3.83 to 2.37pg/insect) but also completely blocked the ability of glucose to elevate JH III (Figure 4G). Furthermore, knockdown of *TOR* downregulated *JHAMT*, *Kr-h1*, and *Met* by 59.87%, 56.20%, and 50.72%, respectively, without affecting *JHE* or *JHEH* (Figure 4H), demonstrating that glucose requires TOR to link to the JH pathway.

**Figure 4.**
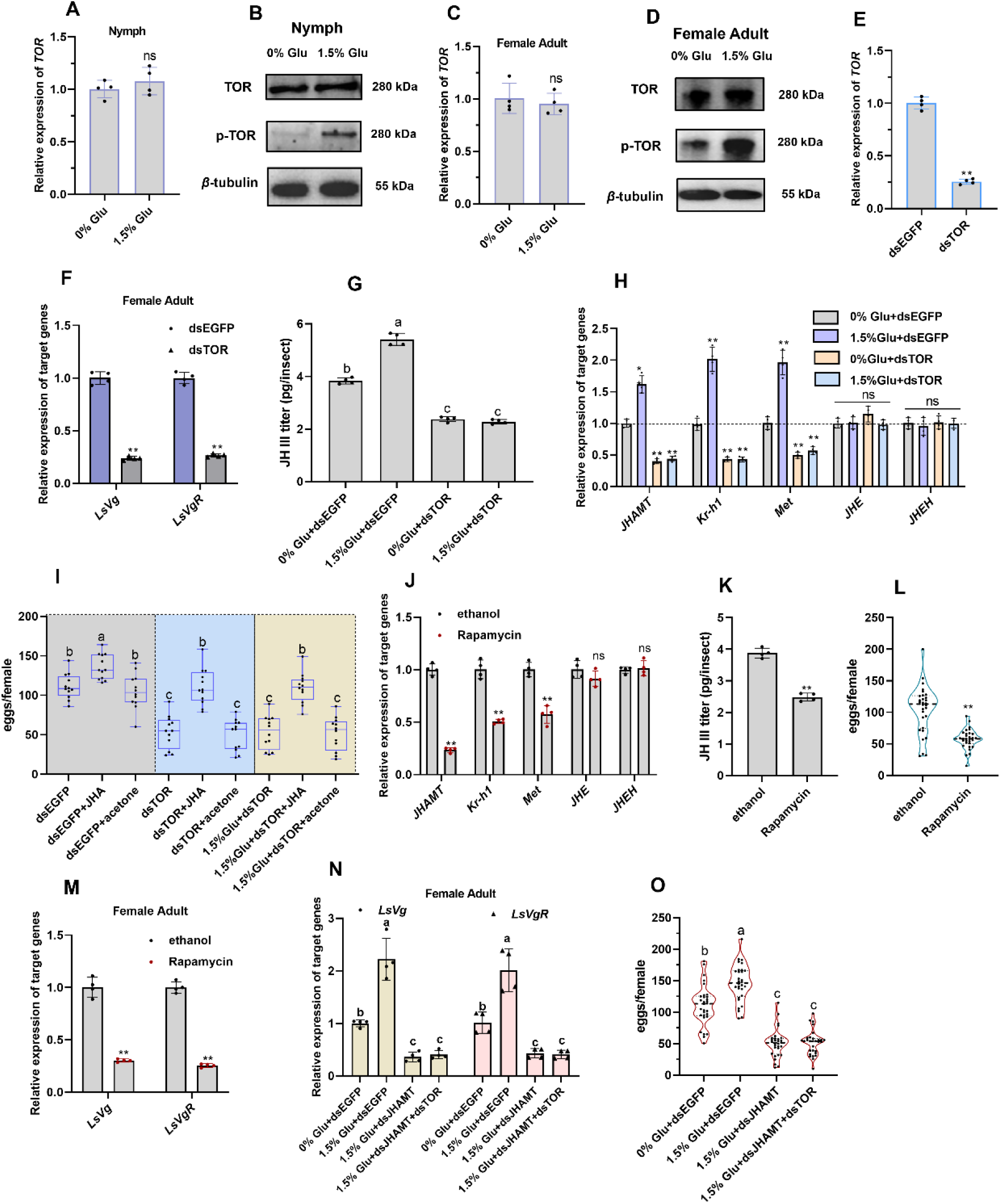
Glucose levels induced by SBPH infestation regulate fecundity through the TOR– JH-Vg pathway. (A, B, C, D) Relative mRNA expression, protein abundance, and phosphorylation levels (Ser2448) of TOR in nymphs and females. (A, B) Third-instar nymphs were maintained for 5 days or (C, D) until adulthood on rice plants grown in 1.5% glucose-supplemented IRRI solution. Control groups were reared on plants in standard IRRI solution (25 insects/group). (E) Relative *TOR* expression at 5 days post-injection in third-instar nymphs injected with dsTOR and reared on rice in standard IRRI solution; dsEGFP served as control (25 insects/group). (F) Expression of *LsVg* and *LsVgR* in females after injection of third-instar nymphs with dsTOR and rearing on rice in standard IRRI solution; dsEGFP was used as control (25 insects/group). (G, H) JH III titer and relative expression of JH-related genes (*JHAMT*, *Kr-h1*, *Met*, *JHE*, *JHEH*) in nymphs at 5 days post-injection with dsTOR and reared on rice cultured in IRRI solution supplemented with 1.5% or 0% glucose; dsEGFP-injected nymphs served as control (25 insects/group). (I) A factorial design was employed to assess fecundity. Third-instar nymphs were first injected with dsTOR and, after 24 hours, subjected to one of three treatments: JHA application, acetone (vehicle control), or no further treatment. These groups were then reared on rice plants supplemented with either 0% or 1.5% glucose. The control cohort received dsEGFP injections followed by the same JHA, acetone, or no-treatment regimens, and was reared exclusively on 0% glucose-supplemented rice. After eclosion, adults were paired (1♀:1♂ per group) and continued to be reared on correspondingly treated plants. Fecundity was measured as the total number of eggs laid per female. (J–M) Gene expression, JH III titer, and oviposition rate following rapamycin treatment. Third-instar nymphs were injected with rapamycin or ethanol (vehicle control) and reared on rice in standard IRRI solution. (J) mRNA levels of *JHAMT*, *Kr-h1*, *Met*, *JHE*, and *JHEH* in nymphs at 5 days post-injection (25 insects/group). (K) JH III titer in nymphs at 5 days post-injection (25 insects/group). (L) Oviposition rate (1♀:1♂ per group). (M) Expression of *LsVg* and *LsVgR* in females (25 insects/group). (N) Expression of *LsVg* and *LsVgR* in females after injection at the third-instar stage with dsEGFP, dsJHAMT, or a mixture of dsTOR and dsJHAMT, and reared on rice plants grown in 1.5% glucose-supplemented IRRI solution (25 nymphs/group; n = 4). Control nymphs were injected with dsEGFP and reared on plants in standard IRRI solution. (O) Oviposition rate of the treated insects in (N) (1♀:1♂ per group). Panel images (A, C, E, F, G, H, J, K, M, N) represent data from four biological replicates, while (B, D), (I), and (L, O) include three, twelve, and thirty replicates, respectively. Data are presented as mean ± s.d. (A, C, E, F, H, J-M) Were analyzed by Student’s *t*-test (**\****p* < 0.05, **\*\****p* < 0.01, ns = not significant). (G, I, N, O) Significant differences (one-way ANOVA with Tukey’s test; *p* < 0.05) are indicated by lowercase letters. JHA, juvenile hormone analog; Glu, Glucose.

The functional hierarchy was confirmed by rescue experiments. Methoprene (a juvenile hormone analog, JHA) application increased egg production by 29.52% (from 111.08 to 136.00 eggs/female; Figure 4I) in the dsEGFP treated group. In addition, treatment with JHA increased the fecundity by 106% (from 53.58 to 110.50 eggs/female) in *TOR*-silenced insects, fully restoring it to the control level (Figure 4I). Furthermore, under dsTOR treatment, high glucose failed to enhance fecundity (53.25 eggs/female), whereas JHA application completely rescued egg production (109.42 eggs/female) (Figure 4I). These results collectively position glucose upstream of the TOR–JH axis. Consistently, rapamycin-mediated TOR inhibition suppressed *JHAMT*, *Kr-h1*, and *Met* expression by 76.70%, 49.65%, and 42.90%, respectively (Figure 4J), reduced JH III titers by 35.92% (2.48 vs. 3.88 pg/insect; Figure 4K), decreased fecundity by 44.9% (57.5 vs. 104.3 eggs/female; Figure 4L), and downregulated *LsVg/LsVgR* by 69.62–74.95% (Figure 4M). Finally, under 1.5% glucose treatment, *JHAMT* silencing reduced *LsVg* and *LsVgR* by 67.95% and 57.91%, respectively, and decreased fecundity by 54.41% (Figure 4N, 4O). Notably, co-knockdown of *TOR* and *JHAMT* did not further suppress *LsVg/LsVgR* or fecundity compared to *JHAMT* knockdown alone (Figure 4N, 4O), confirming that both genes operate in the same linear pathway. Collectively, these results define a coherent signaling cascade in which host-derived glucose activates TOR kinase, which in turn stimulates JH biosynthesis and signaling to enhance vitellogenin production and ultimately increase SBPH fecundity.

### Glucose levels induced by SBPH infestation enhances imidacloprid tolerance by co-activating GST through metabolic and regulatory pathways

Exposure to the LC₅₀ of imidacloprid (2.26 mg/L) for 5 days significantly increased GST activity in SBPH nymphs by 2.04-fold (Figure 5A), and chemical inhibition of GSTs with diethyl maleate (DEM) increased insect mortality by 33.5% under imidacloprid stress (Figure 5B), confirming GSTs’ role in imidacloprid detoxification. Furthermore, feeding on glucose-supplemented (0-2.0% in IRRI) rice enhanced GST activity in a dose-dependent manner (1.23–1.51 fold; Figure 5C), suggesting that glucose-induced tolerance (Tables S1, S2) is mediated, at least in part, by this detoxification system. We next explored how glucose regulates GST activity. Glucose significantly upregulated the expression of *GCL*, the rate-limiting enzyme in GSH synthesis, by 1.47- to 2.21-fold (Figure 5D). This was accompanied by increased cellular GSH content (Figure 5E) and elevated GST activity (Figure 5C). RNAi-mediated knockdown of *GCL* (48.67% reduction; Supplementary Figure S1) reduced both GSH levels by 33.46% and GST activity by 51.77% (Figures 5F, 5G), confirming that glucose enhances GST activity through the GCL–GSH pathway. However, glucose supplementation only partially restored GST activity in *GCL*-silenced insects (Figure 5G), and mortality under imidacloprid (LC_50_ concentration; 2.26 mg/L) exposure remained at an intermediate level (59.5%), between *GCL*-silenced (68%) and glucose-supplemented dsEGFP controls (34%) (Figure 5H). These results indicate that, beyond metabolic control of GSH synthesis, an additional glucose-dependent pathway contributes to GST regulation.

**Figure 5.**
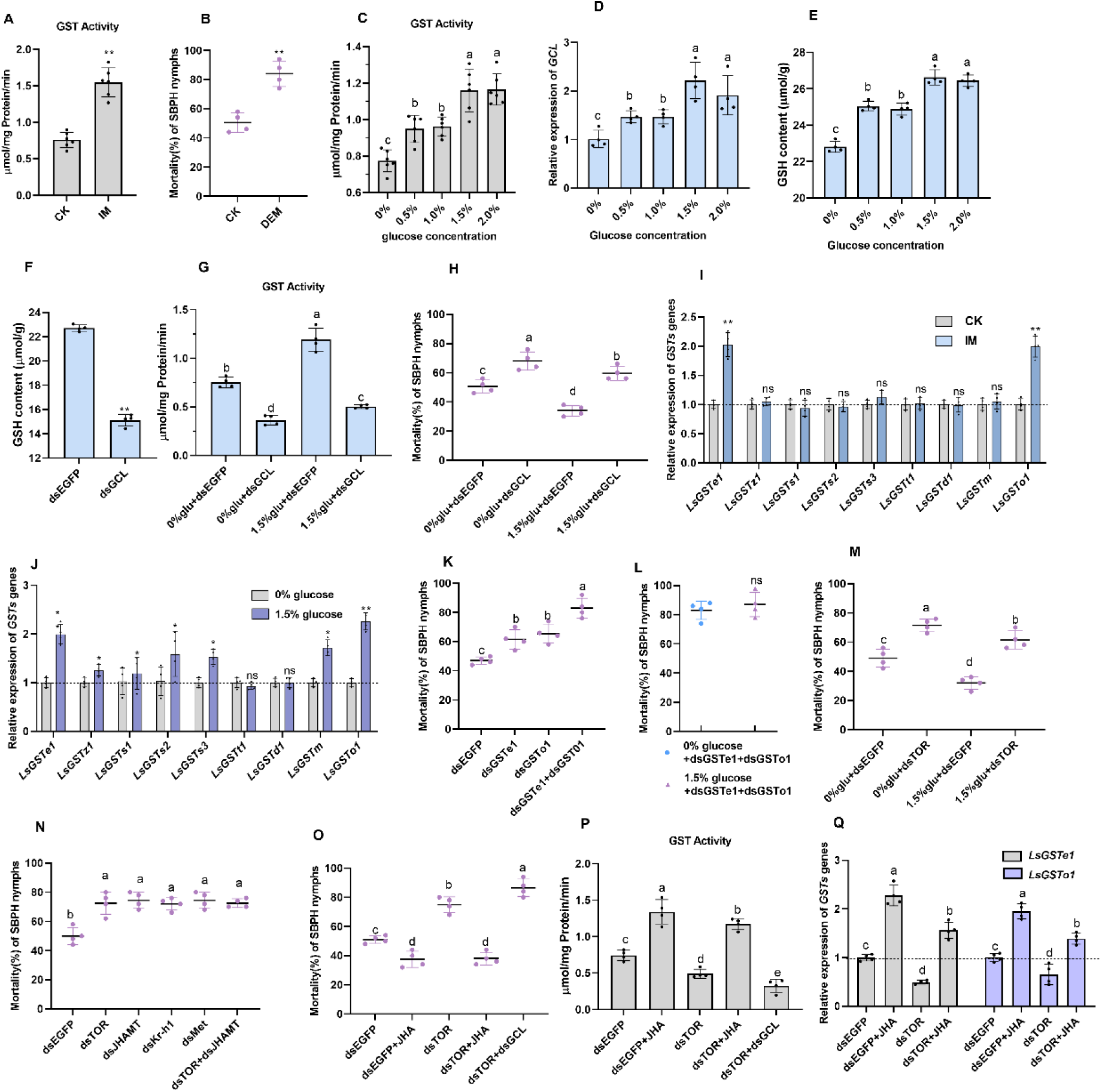
Glucose levels induced by SBPH infestation influence imidacloprid tolerance in SBPH by regulating GST activity. (A) GST activity in nymphs treated with the LC₅₀ dose of imidacloprid (2.26 mg/L) as third-instars and detected 5 days later. Control nymphs were treated with an imidacloprid-free solution under identical conditions (25 nymphs/group). (B) Nymph mortality treated with a combination of imidacloprid and the GST inhibitor DEM as third-instars and recorded at 5 days post-treatment (50 nymphs/group). (C) GST activity in nymphs reared on rice plants supplemented with 0–2.0% glucose as third-instars and assayed 5 days later (25 nymphs/group). (D) *GCL* expression and (E)GSH content in nymphs reared on rice supplemented with 0–2.0% glucose as third-instars and assayed 5 days later (25 insects/group). (F) GSH content in nymphs injected with ds*GCL* as third-instars and assayed 5 days later (25 insects/group). (G) GST activity of nymphs injected with ds*GCL* as third-instars, then reared on rice seedlings cultured in 0% or 1.5% glucose IRRI solution and assayed 5 days later; dsEGFP-injected nymphs served as controls (25 nymphs/group). (H) Mortality rate of insects treated like (G) exposure with LC_50_ imidacloprid for 5 days (50 nymphs/group). (I) Relative expression of 9 *GST* genes in nymphs treated with the LC₅₀ dose of imidacloprid as third-instars and detected 5 days later (25 nymphs/group). (J) Relative expression of 9 *GST* genes in nymphs reared on rice plants supplemented with 0% or 1.5% glucose as third-instars and assayed 5 days later (25 nymphs/group). (K) Mortality of nymphs injected with dsRNA targeting *GSTe1*, *GSTo1*, or both as third-instars, then exposed to the LC₅₀ dose of imidacloprid and recorded 5 days later (50 nymphs/group). (L) Mortality of nymphs with co-knockdown of *GSTe1* and *GSTo1* treated with the LC₅₀ dose of imidacloprid as third-instars, then reared on rice irrigated with 0% or 1.5% glucose solution and recorded 5 days later (50 nymphs/group). (M) Mortality of nymphs subjected to RNAi-mediated *TOR* knockdown as third-instars, then exposed to the LC₅₀ dose of imidacloprid and reared for 5 days on rice cultured in IRRI solution supplemented with 0% or 1.5% glucose; dsEGFP-injected nymphs served as the control (50 nymphs/group). (N) Mortality of nymphs at 5 days post-treatment with the LC₅₀ dose of imidacloprid, following RNAi-mediated knockdown of *TOR*, *JHAMT*, *Kr-h1*, *Met*, combined *TOR* and *JHAMT,* initiated at the third-instar stage (50 nymphs/group). (O) Mortality of nymphs, treated with the LC₅₀ of imidacloprid as third-instars and assayed 5 days later, following RNAi-mediated *TOR* knockdown; *TOR* knockdown with JHA rescue; or combined *TOR* and *GCL* knockdown; compared to the dsEGFP-injected control (50 nymphs/group). (P) GST activity in nymphs 5 days after undergoing RNAi-mediated *TOR* knockdown; *TOR* knockdown with JHA rescue; or combined *TOR* and *GCL* knockdown, compared to the dsEGFP-injected control (25 nymphs/group). (Q) Relative expression of *LsGSTe1* and *LsGSTo1* in nymphs 5 days after RNAi-mediated *TOR* knockdown or *TOR* knockdown with JHA rescue, compared to the dsEGFP-injected control (25 nymphs/group). Panel images (A, C) represent data from six biological replicates, while (B, D, E, F, G, H, I, J, K, L, M, N, O, P, Q) include four replicates. Data are presented as mean ± s.d. (A, B, F, I, J, L) Were analyzed by Student’s *t*-test (**\****p* < 0.05, **\*\****p* < 0.01, ns = not significant). (C, D, E, G, H, K, M-Q) Significant differences (one-way ANOVA with Tukey’s test; *p* < 0.05) are indicated by lowercase letters. IM, imidacloprid; DEM, diethyl maleate; JHA, juvenile hormone analog; Glu, Glucose.

Given the relationship between glucose and the TOR–JH pathway, we next investigated whether this axis influences SBPH tolerance to imidacloprid. We first screened 9 GST genes previously identified in SBPH^44^ and found that only *LsGSTe1* (epsilon class) and *LsGSTo1* (omega class) were significantly induced by the LC₅₀ of imidacloprid (2.02- and 1.97-fold; Figure 5I). Glucose similarly upregulated these two genes (1.98- and 2.25-fold), along with several other *GSTs* (1.16–1.71 fold), except *LsGSTt1* and *LsGSTd1* (Figure 5J). Functional validation showed that RNAi knockdown of *LsGSTe1* or *LsGSTo1* (51.06% and 50.57% reduction, respectively; Supplementary FigureS1) increased nymph mortality by 14.5% and 18.5%, while co-knockdown (*LsGSTe1* and *LsGSTo1* reduced by 72.08% and 53.91%, respectively; Supplementary Figure S1) increased mortality by 36% under imidacloprid exposure (Figure 5K). Importantly, glucose supplementation failed to enhance tolerance in co-silenced nymphs (87% vs. 83% mortality; Figure 5L), demonstrating that both *LsGSTe1* and *LsGSTo1*are required for glucose-induced tolerance.

We next tested the involvement of TOR–JH pathway in imidacloprid tolerance as well as GST regulation. We found that silencing *TOR* significantly increased nymph mortality under LC₅₀ imidacloprid treatment (from 49.0% to 71.5%), and glucose supplementation only partially rescued this phenotype (to 61.5%), suggesting that *TOR* positively regulates tolerance (Figure 5M). Similarly, separate knockdown of JH-related genes (*JHAMT*, *Kr-h1*, or *Met*) increased mortality to 72–74.5% (Figure 5N). In contrast, JHA supplementation enhanced tolerance in dsEGFP controls (from 51% to 37.5% mortality; Figure 5O), increased GST activity by 1.8-fold (Figure 5P), along with upregulation of *LsGSTe1* and *LsGSTo1* (2.28- and 1.95-fold; Figure 5Q), establishing JH as a positive regulator of GST-mediated detoxification. Additionally, *TOR* knockdown reduced GST activity by 50.97% (Figure 5P) and downregulated *LsGSTe1* and *LsGSTo1* expression by 50.19% and 35.08%, respectively (Figure 5Q), establishing TOR as a positive regulator of GST-mediated detoxification. The absence of additive mortality in *TOR*/*JHAMT* co-silencing (Figure 5N), together with the complete rescue by JHA in *TOR*-silenced insects, demonstrates that TOR acts upstream of JH to regulate detoxification. Upon JHA rescue, imidacloprid-induced mortality decreased from 75% to 38% (Figure 5O), GST activity increased by 3.21-fold (Figure 5P), and the expression levels of *LsGSTe1* and *LsGSTo1* were upregulated by 3.16-fold and 2.13-fold, respectively (Figure 5Q), compared with the dsTOR-only group.

Finally, to integrate both regulatory and metabolic axes, we generated *TOR* and *GCL* double-knockdown insects. This combination resulted in the highest mortality (86.5%) under imidacloprid exposure, substantially greater than that of either *TOR*-silenced (75.01%) or *GCL*-silenced (68.01%) insects (Figures 5O, 5H). Correspondingly, GST activity was most pronouncedly suppressed in the co-knockdown group, showing a 56.27% reduction compared to *TOR* silencing alone and a 10.50% reduction compared to *GCL* silencing alone (Figures 5P, 5G). These additive effects indicate that TOR–JH signaling and GCL–GSH metabolism act cooperatively to modulate GST activity, jointly determining imidacloprid tolerance in SBPH.

## Discussion

Our study reveals a conserved nutrient-responsive mechanism whereby SBPH infestation elicits a carbohydrate metabolism shift in rice, and the insect subsequently utilizes host-derived glucose to simultaneously augment its reproductive capacity and insecticide tolerance. Specifically, we identify host-derived glucose as a central signaling molecule that modulates two interconnected molecular cascades exerting dual fitness benefits: a glucose-TOR-JH-Vg signaling axis governing fecundity, and a dual metabolic–regulatory pathway that coordinates GST activity to confer imidacloprid tolerance. Importantly, multiple lines of evidence indicate that glucose functions here as a regulatory signal rather than merely a caloric resource, as its effects depend on TOR phosphorylation and endocrine activation and cannot be mimicked by osmotic controls.

This phenomenon aligns with the classic paradigm in plant-insect interactions in which herbivores subvert plant defensive or compensatory metabolic responses for their own benefit^9,45,46^. Consistent with earlier reports that planthoppers manipulate host sugar allocation through salivary effectors^9^, our results demonstrate that SBPH infestation induces a systemic, time- and density-dependent accumulation of glucose in rice aerial tissues. Notably, this response displays a biphasic pattern, with glucose levels peaking at moderate infestation densities and declining under extreme pest pressure, suggesting the existence of a physiological threshold beyond which plant regulatory capacity becomes constrained. Such a non-linear response argues against an artefactual effect driven by excessive manipulation and instead points to inherent physiological limits in plant metabolic regulation under herbivore pressure. Strikingly, this metabolic shift is further amplified by gravid females, revealing that oviposition behavior exerts a distinct and additive influence on host carbohydrate metabolism beyond that induced by feeding alone. Notably, the upstream signals (e.g., specific salivary effectors secreted by SBPH or oviposition-associated plant response factors) that trigger glucose reallocation in rice remain uncharacterized and represent a key direction for future in-depth research.

Crucially, through a series of complementary experiments, we established glucose as the specific mediator of enhanced SBPH fitness. Increased fecundity and upregulation of *LsVg*/*LsVgR* were induced by metabolizable sugars (glucose, sucrose, and trehalose), but not by the non-metabolizable osmotic control mannitol. Given that osmotic pressure, a key determinant of plant cell turgor pressure, can disrupt insect homeostasis and impair fitness when insects ingest hyperosmotic plant sap^47,48^, we rigorously excluded confounding effects of rice osmotic pressure in this study. Importantly, inhibition of sucrase or trehalase abolished the reproductive benefits of sucrose and trehalose, and these effects were fully restored by exogenous glucose supplementation. This coherent series of loss- and rescue-of-function experiments conclusively rules out osmotic effects and establishes glucose as the active nutritional signal. Together, these results distinguish glucose-specific signaling effects from general carbon availability, reinforcing the view that glucose acts upstream of defined physiological pathways rather than simply enhancing energetic status. Beyond reproduction, host-derived glucose significantly bolstered SBPH tolerance to imidacloprid. The elevated LC₅₀ values on pre-infested or glucose-supplemented plants, and the glucose-specific nature of this tolerance (mimicked by metabolizable disaccharides and negated by their hydrolase inhibitors), demonstrate a direct link between nutritional status and detoxification capacity. Notably, studies have shown that brown planthopper (*Nilaparvata lugens*) infestation can reshape sugar distribution in rice by altering the expression of rice sugar transporters, yet the mechanism through which planthoppers regulate these transporters remains unresolved^9^. Given that elevated sugar levels might enhance plant stress resistance^49,50^, our study reveals an intriguing ecological paradox that SBPH infestation likely manipulates glucose distribution via unidentified pathways to boost its own fitness and resilience. The biphasic glucose response further implies that plant metabolic defenses may be optimized only within a limited infestation range, a feature that could contribute to the nonlinear dynamics of SBPH population outbreaks. In summary, SBPH orchestrates a profound manipulation of host rice, systemically altering glucose distribution and levels. The insect then exploits on this manipulated nutritional landscape, deriving dual benefits of increased fecundity and enhanced insecticide tolerance. This strategy underscores the evolutionary sophistication of herbivorous insects in turning host plant responses to their advantage.

A key finding of this work is the discovery of a glucose-TOR-JH-Vg signaling axis that underlies the enhanced fecundity in SBPH. While the insect TOR kinase is well-established as a nutrient sensor, and its activation typically responds to amino acid availability via phosphorylation at Ser2448 to govern growth and reproduction^51–53^, sugar-promoted TOR activation has been reported in *Drosophila*²⁹, and our study extends this conserved regulatory mechanism to hemipteran insects. In this study, we show that host-derived glucose induces TOR phosphorylation at Ser2448 in SBPH, a conserved mechanism mirroring mechanisms in mammals and plants^26–28^. Expanding upon the amino acid-TOR-JHAMT axis reported in other insects^23–25^, our results revealed that glucose-mediated TOR activation acts as the critical upstream trigger that stimulates the JH pathway to enhance fecundity in SBPH. Further RNAi-and pharmacology-based analyses establish the functional indispensability and hierarchical organization of this pathway. *TOR* knockdown or rapamycin inhibition suppressed *JHAMT*, *Kr-h1*, and *Met* expression, reduced JH III titers, and markedly impaired vitellogenin expression and egg production. Epistasis analysis further resolved the genetic order of this cascade: JHA fully rescued the fecundity defect in *TOR*-silenced insects, whereas simultaneous knockdown of *TOR* and *JHAMT* produced no additive effect compared with *JHAMT* silencing alone. It should be noted that while absolute JH titers were quantified using ELISA, this method is both widely adopted and well-established in the field and the conclusions are strongly supported by concordant transcriptional responses and functional rescue experiments, which establish the role of JH signaling in glucose-induced fecundity. These findings place glucose upstream of TOR-JH pathway. Collectively, our results define a coherent glucose-TOR-JH-Vg axis through which host nutritional status is transduced into endocrine control of SBPH reproduction. This nutrient-responsive cascade might be conserved across insect species.

Parallel to fecundity enhancement, SBPH utilizes host glucose to bolster its tolerance to the insecticide imidacloprid by supporting a novel dual-pathway model for GST activation, entailing both metabolic fueling and transcriptional regulation. GSTs are well-established contributors to insecticide resistance through xenobiotic detoxification and antioxidant defense^54,55^, and increased GST activity has been repeatedly associated with neonicotinoid resistance in planthoppers^56–59^. Our data demonstrate that glucose-induced tolerance is mediated by enhanced GST activity operating through two cooperative mechanisms. The first is a metabolic axis in which glucose upregulates *GCL*, the rate-limiting enzyme for GSH biosynthesis, thereby increasing the availability of the essential GST co-substrate. Consistent with this model, glucose elevated *GCL* expression, increased GSH content, and enhanced GST activity, whereas *GCL* knockdown markedly reduced both GSH levels and GST activity and compromised glucose-induced tolerance. The second is a regulatory axis in which glucose activates *GSTs* gene expression through the TOR–JH signaling cascade. Among the nine *GSTs* genes examined, *LsGSTe1* and *LsGSTo1* emerged as key effectors jointly induced by imidacloprid and glucose. Functional analyses confirmed their necessity for detoxification, as individual and combined silencing of these genes significantly increased insect mortality and abolished glucose-mediated tolerance. Notably, JHA application rescued the GST activity and *LsGSTe1/LsGSTo1* expression suppressed by *TOR* knockdown, positioning TOR upstream of JH in the regulation of GST-mediated detoxification. Most compellingly, simultaneous knockdown of *TOR* and *GCL* produced the highest mortality and the strongest suppression of GST activity, demonstrating that the TOR-JH regulatory axis and the GCL-GSH metabolic pathway act non-redundantly and cooperatively to potentiate GST activity and determine imidacloprid tolerance. Beyond the GCL-GSH-GST and TOR-JH-GST pathways characterized in this study, elevated glucose may also enhance insecticide detoxification through additional routes (e.g., providing carbon skeletons for phase II xenobiotic conjugation reactions or fueling energy-dependent detoxification processes in insect midgut and fat body), which warrant further experimental verification.

The identification of the glucose-TOR-JH axis as a key regulator of SBPH fecundity and insecticide tolerance provides novel strategies for eco-friendly, nutrient-based pest control. Firstly, varieties that limit SBPH-induced glucose redistribution would reduce reproduction and insecticide tolerance without yield loss. Secondly, small-molecule inhibitors targeting TOR phosphorylation or JH synthesis can serve as biopesticides or synergists to improve insecticide efficacy and delay resistance. Finally, optimized fertilization and irrigation can reduce shoot glucose accumulation and suppress SBPH outbreaks. These strategies offer sustainable alternatives to traditional insecticides and help mitigate insecticide resistance of SBPH.

In conclusion, our research unveils a multi-faceted manipulation where SBPH infestation triggers a systemic metabolic shift in rice, leading to elevated glucose in the aerial phloem. The insect then exploits this enriched resource to simultaneously power a reproductive program via the glucose-TOR-JH-Vg cascade and fortify its detoxification system via coordinated GCL-GSH (metabolic) and TOR-JH-GST (regulatory) pathways. This work profoundly expands our understanding of nutrient-mediated plant-insect interactions by positioning glucose not merely as a food source, but as a critical signaling molecule that an herbivore harnesses to optimize its fitness. These insights into the sophisticated resource-manipulation tactics of SBPH pave the way for innovative pest control strategies that target these specific nutrient-sensing and detoxification interfaces.

### Limitations and Future Research Directions

This study focuses exclusively on SBPH and the neonicotinoid insecticide imidacloprid, and thus the generalizability of the observed glucose-mediated fitness enhancement remains to be validated. Future research will focus on three key directions: (1) Verifying the conservation of the glucose-TOR-JH axis in other economically important rice planthoppers; (2) Expanding insecticide tolerance assays to other commonly used rice insecticides (e.g., thiamethoxam, pymetrozine, triflumezopyrim) to determine whether the glucose-mediated detoxification pathway confers broad-spectrum tolerance; (3) Identifying the specific SBPH salivary effectors and plant signaling pathways that trigger glucose reallocation in rice, to complete the mechanistic framework of host metabolic changes manipulated by herbivores.

## Materials and Methods

### Insect and Plant Materials

SBPH were collected from paddy fields in Yangzhou, Jiangsu Province, China, and maintained on rice seedlings in a climate-controlled chamber at 26 ± 1°C, 70–80% relative humidity, with a 16-h light/8-h dark photoperiod. Synchronized third-instar nymphs were used for all experiments unless otherwise stated. Ten-day-old rice seedlings of uniform size, cultivated in IRRI (https://www.irri.org/) nutrient solution, were selected for all treatments.

### Plant Infestation Assays

To investigate the effect of SBPH infestation on rice carbohydrate metabolism, the following assays were conducted. (i) Time-course assay: Rice plants were infested with 25 nymphs/plant for 1, 3, and 5 days, then the aerial tissues were collected for glucose measurement. (ii) Density-dependent assay: Plants were infested with 0, 1, 5, 10, 15, 20, 25, or 30 nymphs/plant for 5 days, after which aerial tissues were collected for glucose quantification. (iii) Life stage- and sex-dependent assay: Plants were infested with 25 insects/plant representing distinct developmental stages or sexes (nymphs, virgin females, gravid females, or males) for 5 days, then the aerial tissues were collected for glucose measurement and gene expression analysis. (iv) Systemic response assay: After a 5-day infestation with 25 third-instar nymphs, root glucose, whole-plant glucose levels were measured. All experiments included 4 replicates.

### Pre-Infestation assays

For the pre-infestation assay, rice seedlings were pre-infested with 25 nymphs/plant for 3 days or kept uninfested. After removing the initial nymphs, fresh nymphs (25/plant) were introduced. Biochemical and molecular analyses were performed on test nymphs at 5 days post-infestation and on newly emerged female adults (within 24 h post-eclosion). For nymphal samples, we quantified glucose levels, JH III titers, and the mRNA expression of key JH pathway genes (including *JHAMT*, *Kr-h1*, *Met, JHE*, and *JHEH*). For newly emerged females, we measured glucose levels and analyzed the mRNA expression of *vitellogenin gene LsVg and its receptor LsVgR*. All these measurements were conducted with four replicates. For fecundity assessment, newly emerged adults were paired (1♀:1♂), and the number of eggs laid was recorded and each treatment included 12 females.

### Carbohydrate Supplementation

For carbohydrate supplementation, rice plants were grown in IRRI nutrient solution supplemented with glucose (G7021, Sigma-Aldrich, US) at concentrations of 0.5%, 1.0%, 1.5%, or 2.0% (w/v), and nymphs (25 per plant) were reared on these treated plants, with the culture solution being renewed daily^60^. The mRNA expression of *LsVg* and *LsVgR* was analyzed in newly emerged females, with four replicates. For fecundity assessment, newly emerged adults were paired (1♀:1♂), and the number of eggs laid was recorded, while each treatment including 30 females. Glucose levels were measured in the aerial tissues of plants grown in either 1.5% glucose-supplemented or standard IRRI solution for 5 days, as well as in the nymphs that had fed on these plants for the same duration. Concurrently, the mRNA expression of key JH pathway genes (including *JHAMT*, *Kr-h1*, *Met*, *JHE*, and *JHEH*) was analyzed in nymphs. These experiments included four replicates. To validate the effects of different sugars and osmotic pressure, sucrose (V900116, Sigma-Aldrich), trehalose (T9531, Sigma-Aldrich), and mannitol (M4125, Sigma-Aldrich) solutions isotonic with 1.5% glucose (83.3 mM) in IRRI nutrient solution were applied to rice plants via root irrigation. Mannitol was used as an osmotic control since it is not absorbed by rice plants nor does it participate in their physiological activities. Nymphs (25/plant) were reared on these treated plants. For fecundity assessment, newly emerged adults were paired (1♀:1♂), and the number of eggs laid was recorded, while each treatment including 12 females. Glucose levels were quantified in nymphs at 5 days post-infestation and the experiments included 4 replicates.

### Exogenous Glucose and Hydrolase Inhibitor Injection

To elevate glucose levels, each nymph was injected (using a Nanoliter 2010 injector; WPI, USA) with 0.1 μL of a sterile 0.6 mM glucose solution, following a method adapted from Lin et al. (2018)^57^. Control nymphs received an equal volume of water. To inhibit sugar metabolism, nymphs feeding on sucrose- or trehalose-treated rice plants were injected with a sucrase inhibitor (5 mM arabinose (W325501, Sigma-Aldrich) solution, 0.1 μL/nymph) or a trehalase inhibitor (0.2 mM Validamycin A (Val A; MS0048, Maokangbio, CN) solution, 0.1 μL/nymph), respectively. The dosages of arabinose and Validamycin A, which were determined by preliminary experiments and a previous report^61^, showed no significant lethal effects on nymphs. A subset of the inhibitor-treated nymphs received a rescue injection of glucose 24 hours later. For all treatments, glucose levels were quantified in nymphs at 5 days post-infestation and the experiments included 4 replicates. For fecundity assessment, newly emerged adults were paired (1♀:1♂), and the number of eggs laid was recorded, and each treatment included 12 females.

### RNA Extraction, cDNA Synthesis, and Quantitative Real-Time PCR (RT-qPCR)

Total RNA was extracted from SBPH whole bodies and rice tissues using TRIzol Reagent (15596026CN, Invitrogen, USA), with quality verified by electrophoresis and quantified using NanoPhotometer N50 (Implen, GER). cDNA was synthesized using HiScript II RT SuperMix (R233-01, Vazyme, CN). Target genes (the names and accession numbers on NCBI listed in Table S3) expression was analyzed by RT-qPCR using specific primers (Table S4) and ChamQ SYBR Master Mix (Q711-02, Vazyme, CN) on a CFX96T system (Bio-Rad), following MIQE guidelines^62^. *ARF* (JF728807) and *UBQ5* (AK061988) served as reference genes for SBPH and rice, respectively^8,43^.The relative expression was calculated by the 2^-ΔΔCt^ method.

### RNA Interference

Double-stranded RNAs (dsRNAs) targeting *TOR* (350 bp), *JHAMT* (356 bp), *Kr-h1* (330 bp), *Met* (330 bp), *GSTe1*(300 bp), *GSTo1*(300 bp), and *GCL* (306 bp) were synthesized via *in vitro* transcription using the TranscriptAid T7 Kit (K0441, Thermo Fisher Scientific, USA) with gene-specific primers (Table S5). dsEGFP was used as the control. Nymphs were microinjected with 0.1 μL of individual dsRNA solutions (1000 ng/μL) using a Nanoliter 2010 injector (WPI, USA). For dual-gene knockdown experiments, the corresponding two dsRNA solutions were mixed at a 1:1 ratio, maintaining the concentration of each dsRNA at 1000 ng/μL in the final mixture, and 0.1 μL of this mixture was injected per nymph.

### Fecundity Assay

Following their respective treatments during the nymphal stage, newly emerged brachypterous adults (the dominant morphotype) were paired (1♀:1♂) and confined on healthy rice seedlings. These seedlings were replaced daily, and the number of eggs laid was counted under a microscope^63,64^.

### Insecticide Tolerance Bioassay

The tolerance of SBPH to imidacloprid was assessed using a rice seedling dip method. A stock solution of imidacloprid (HY-B0838, MCE, US) was first prepared using acetone plus 10% Tween-20 as an emulsifier. This stock was then serially diluted with water to create working solutions at concentrations of 0, 0.5, 1, 2, 4, 8, 16, and 32mg/L^65^. Bioassays were conducted under several pre-treatment conditions to evaluate their effects on insecticide tolerance: (i) Pre-infestation: Nymphs were reared on rice plants that had been either pre-infested with 25 nymphs for 3 days or left uninfested. (ii) Carbohydrate Supplementation: Rice was grown in IRRI nutrient solution supplemented with glucose (0.5%, 1.0%, 1.5%, or 2.0%, w/v) and nymphs were reared on these treated plants. To validate the effects of different sugars and osmotic pressure, sucrose, trehalose, and mannitol solutions isotonic with 1.5% glucose (83.3 mM) in IRRI nutrient solution were applied to rice plants via root irrigation. Nymphs were reared on these treated plants at a density of 25 insects per seedling. Each treatment group consisted of two seedlings, totaling 50 nymphs. The experiments included 3 replicates. Mortality was recorded at 5 days post-infestation, with insects considered dead if unresponsive to gentle prodding. Concentration-mortality data were analyzed by probit analysis to determine LC₅₀ values.

### Pharmacological and Hormonal Treatments

(i) GST Inhibition: Rice seedlings were pretreated by immersion in a 50 mg/L diethyl maleate (DEM; HY-Y1147, MCE, US) solution (formulated in acetone with 0.1% Tween-20) for 2 hours. Following this pretreatment, the seedlings were subsequently exposed to the LC₅₀ concentration of imidacloprid (2.26 mg/L) for 30 seconds^65^. Control seedlings were subjected to the same imidacloprid treatment but without DEM pretreatment. After the treatments, nymphs were reared on these seedlings at a density of 25 insects per plant. Each treatment group consisted of two seedlings (50 nymphs total), and the experiment included four replicates. (ii) TOR Inhibition: For pharmacological inhibition, each nymph received a 50 nL injection of either 1.0 nM rapamycin (HY-K0010, MCE, US) or an equivalent ethanol solvent (control)^25,53,66^. Treated nymphs (25/plant) were maintained on rice seedlings. The mRNA levels of *JHAMT*, *Kr-h1*, *Met*, *JHE*, and *JHEH* as well as JH III titer in nymphs were measured at 5 days post-injection. In addition, the mRNA expression of *LsVg* and *LsVgR* was analyzed in newly emerged female adults within 24 hours post-eclosion. The experiment included 4 replicates. For fecundity assessment, newly emerged adults were paired (1♀:1♂), and the number of eggs laid was recorded. Each treatment included 30 females. (iii) JH Rescue: For the juvenile hormone analog (JHA) rescue assay, a topical application was performed 24 hours after the nymphs had been microinjected with dsTOR or dsEGFP. Each nymph received 100 nL of methoprene (40596-69-8, Sigma-Aldrich; 10 μg/μL in a 1:10 acetone:ddH₂O solution) or an equivalent volume of the vehicle solution alone^67^.

### Biochemical Assays

(i) Glucose Quantification: Tissues were homogenized in extraction buffer, centrifuged at 8,000 × g for 10 min, and the supernatant was analyzed spectrophotometrically at 505 nm using a Multiskan microplate reader (Thermo Fisher Scientific, USA) and a glucose assay kit (BC2500, Solarbio, China), following the manufacturer’s protocols. (ii) GST Activity: GST activity was measured using a commercial assay kit (BC0355, Solarbio, CN) by quantifying CDNB-GSH conjugation kinetics at 340 nm. (iii) GSH Content: For GSH quantification, nymph homogenates were processed using a GSSG/GSH assay kit (S0053, Beyotime, CN). Samples were split for parallel measurement of total GSH (after GSSG reduction) and GSSG (after GSH masking). Both were determined via TNB formation kinetics, with concentrations calculated from standard curves (nmol/g tissue)^68^. (iv) JH III titer: JH III levels in SBPH samples were determined by a double-antibody sandwich enzyme-linked immunosorbent assay (ELISA) using a commercial kit (YX-100800I, Shanghai Qiaoshe Biotechnology, CN), as previously outlined by Gao et al. (2025)^69^.

### Western Blotting

For protein analysis, pooled SBPH samples were homogenized in RIPA buffer containing protease and phosphatase inhibitors. After centrifugation (12,000×g, 20 min, 4°C), proteins were separated by 10% SDS-PAGE and transferred to PVDF membranes. Membranes were blocked with 5% BSA (for phosphoproteins) or 5% nonfat milk in TBS at 4°C overnight. Primary antibodies included Anti-Phospho-mTOR (Ser2448) rabbit polyclonal antibody (1:2000; AF5869, Bryotime); Anti-TOR rabbit polyclonal antibody (1:1000; FNab05417, FineTest, CN); Anti-β-Tubulin mouse monoclonal antibody (1:2000; AF2835, Bryotime). After TBST washes, membranes were incubated with HRP-conjugated secondary antibodies (1:5000; Sigma-Aldrich) for 1 h. Protein bands were visualized using ECL substrate and quantified using Image Lab (Bio-Rad). All antibodies used were validated for specificity (Supplementary Figure S2).

### Statistical Analysis

SPSS software (version 25.0, IBM, Armonk, NY, USA) was utilized for data analysis. Student’s *t*-test was used to assess statistically significant differences between two groups. For multiple group comparisons, statistical analyses were carried out by one-way ANOVA followed by Tukey’s post hoc test. A *p*-value < 0.05 was considered statistically significant.

### Supplemental information

**Table S1.** Effects of different treatments on the resistance to imidacloprid in SBPH.

| treatment | LC<sub>50</sub> (mg/L, 95% CI) | slope |
| --- | --- | --- |
| No Pre-infestation | 2.26 (1.34-3.19) | 1.94±0.24 |
| Pre-infestation | 5.85 (3.68-7.99) | 1.38±0.18 |
| 0.5% glu | 2.35 (1.40-3.31) | 1.96±0.24 |
| 1.0% glu | 5.83 (3.73-7.89) | 1.46±0.19 |
| 1.5% glu | 6.56 (4.29-8.77) | 1.45±0.19 |
| 2.0% glu | 6.46 (4.22-8.58) | 1.50±0.19 |

**Table S2.**
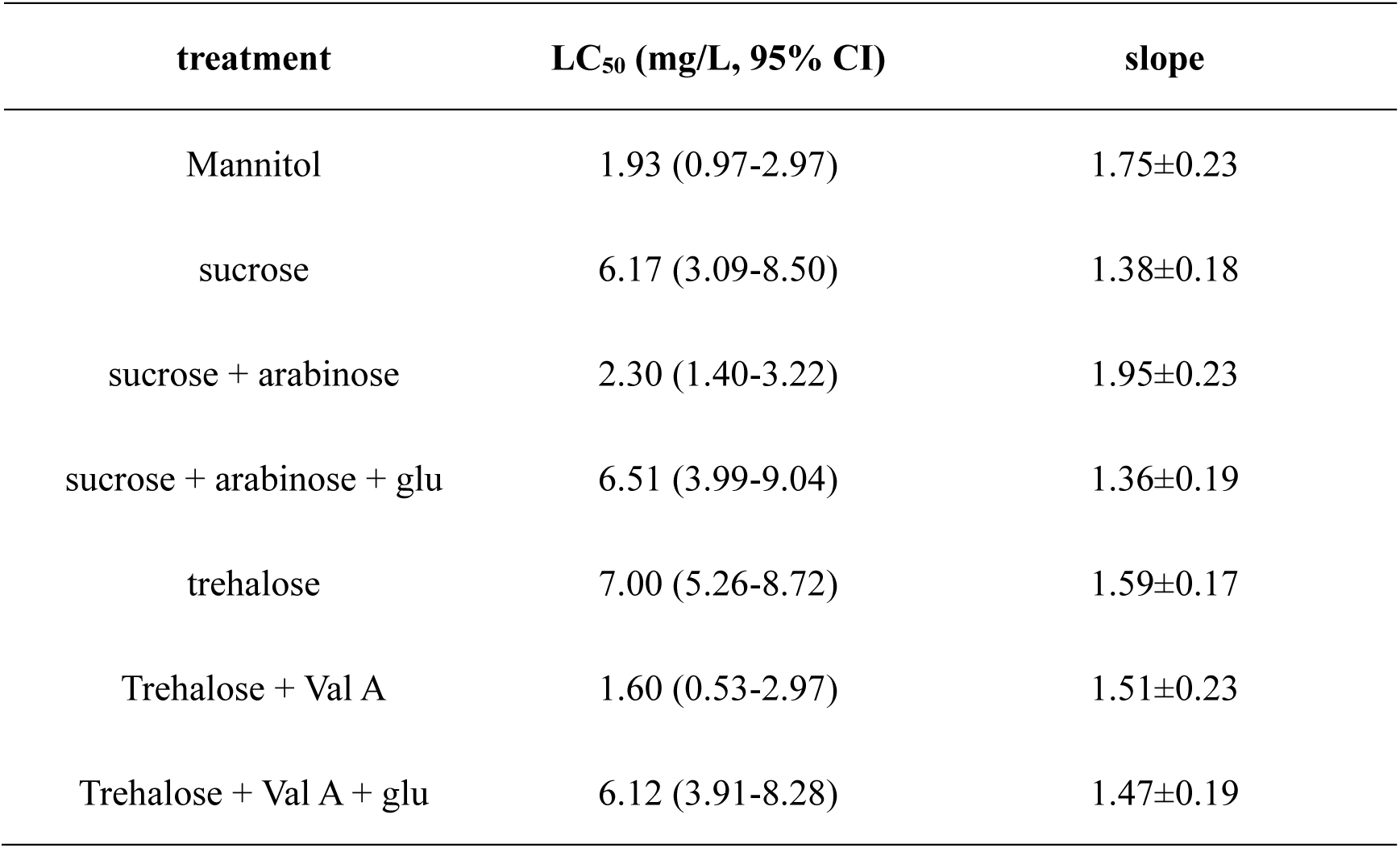
Effects of different sugars on the resistance to imidacloprid in SBPH.

| treatment | LC<sub>50</sub> (mg/L, 95% CI) | slope |
| --- | --- | --- |
| Mannitol | 1.93 (0.97-2.97) | 1.75±0.23 |
| sucrose | 6.17 (3.09-8.50) | 1.38±0.18 |
| sucrose + arabinose | 2.30 (1.40-3.22) | 1.95±0.23 |
| sucrose + arabinose + glu | 6.51 (3.99-9.04) | 1.36±0.19 |
| trehalose | 7.00 (5.26-8.72) | 1.59±0.17 |
| Trehalose + Val A | 1.60 (0.53-2.97) | 1.51±0.23 |
| Trehalose + Val A + glu | 6.12 (3.91-8.28) | 1.47±0.19 |

**Table S3.**
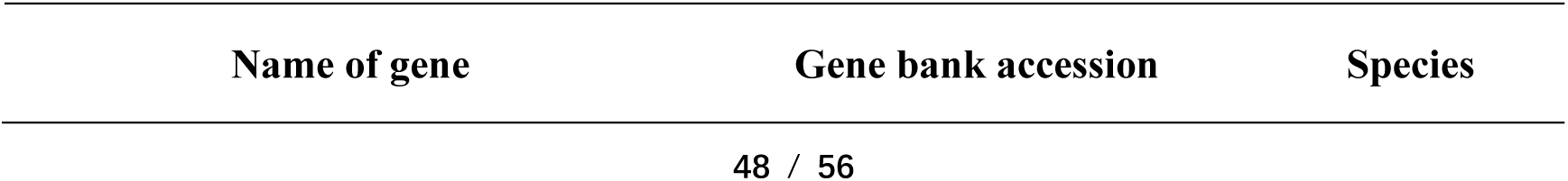
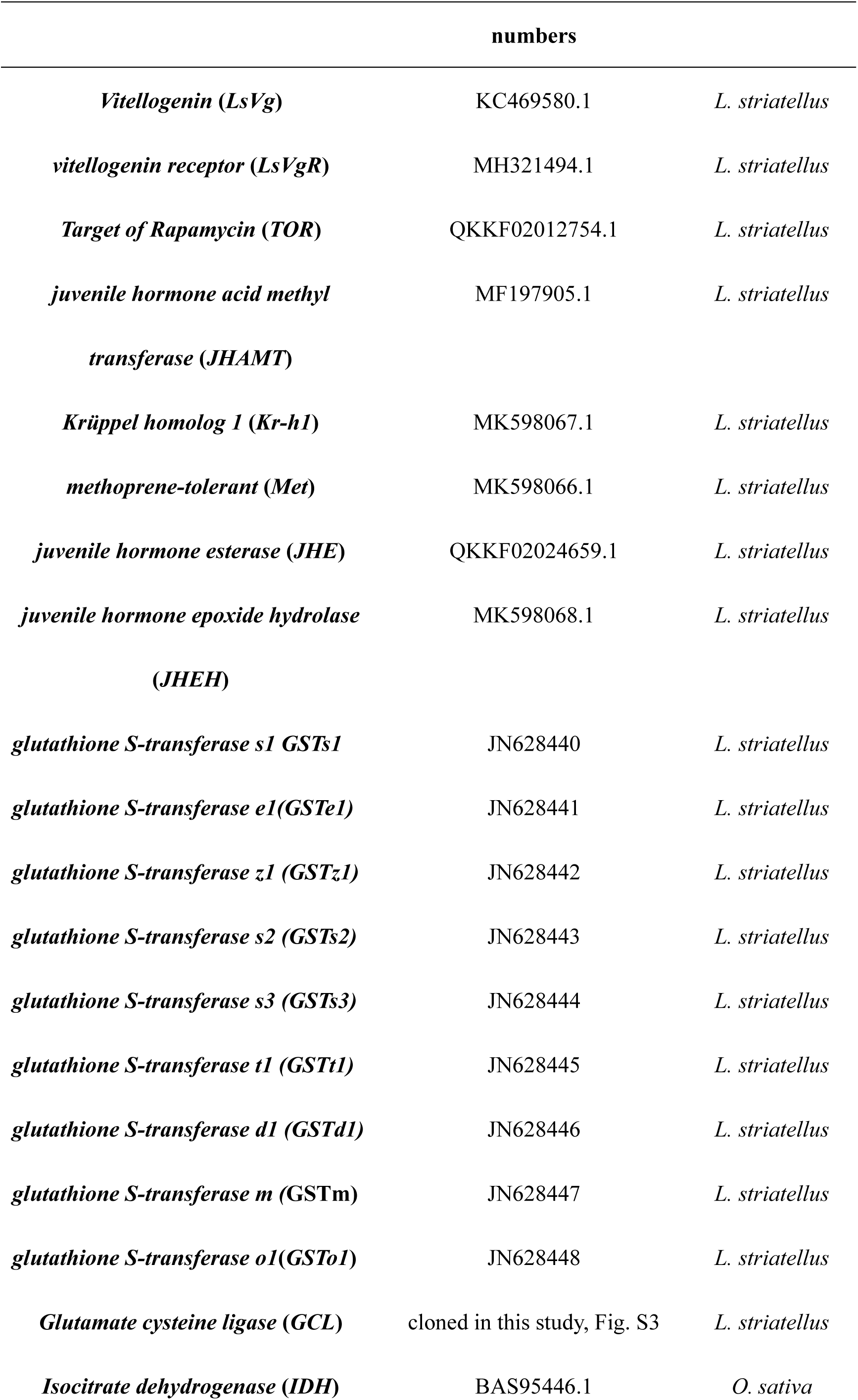
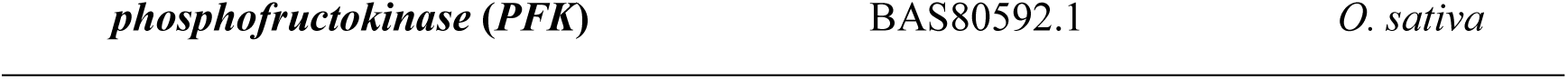
Target genes’ names were detected with RT-qPCR in this study Name of gene Gene bank accession Species.

**Table S4.**
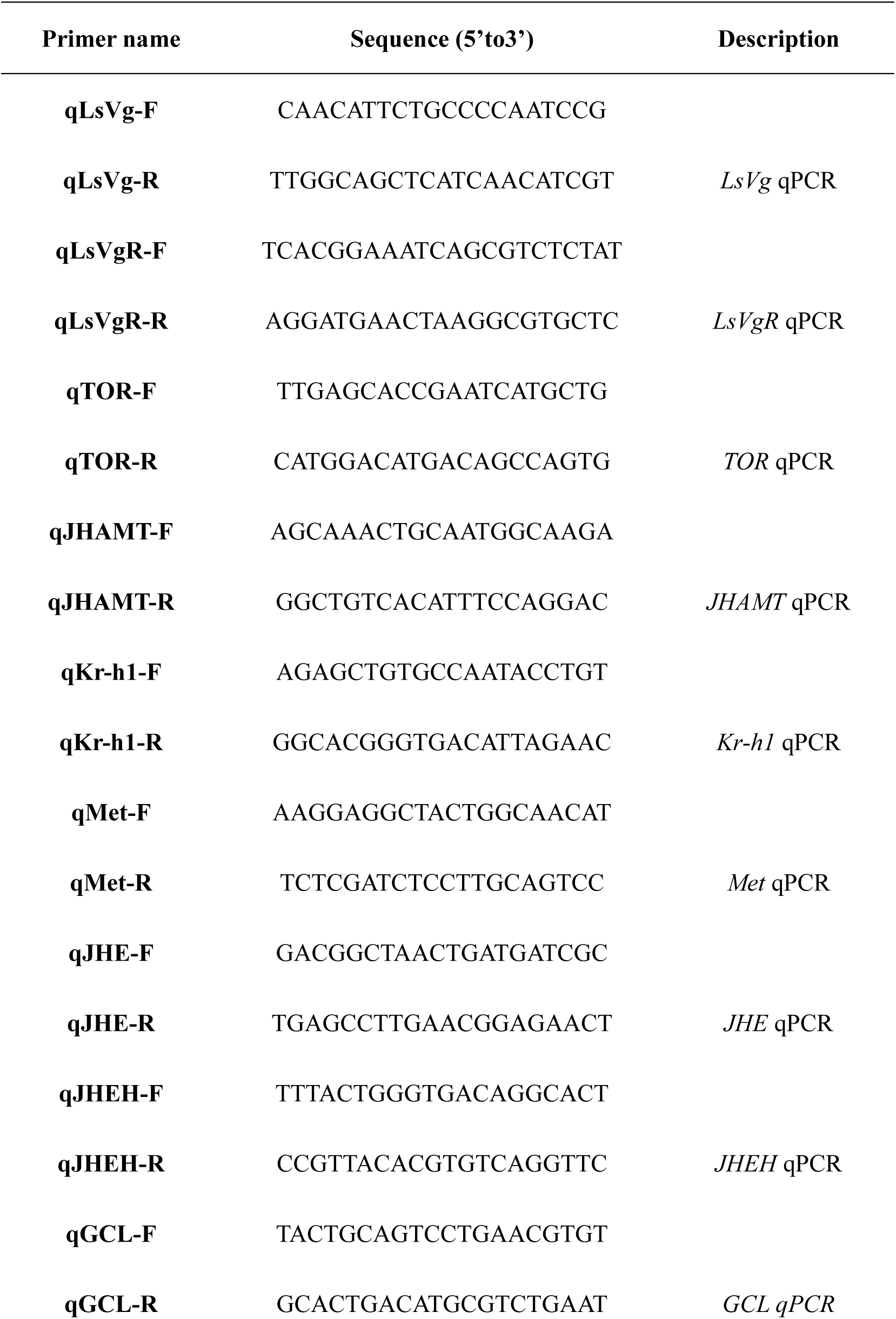
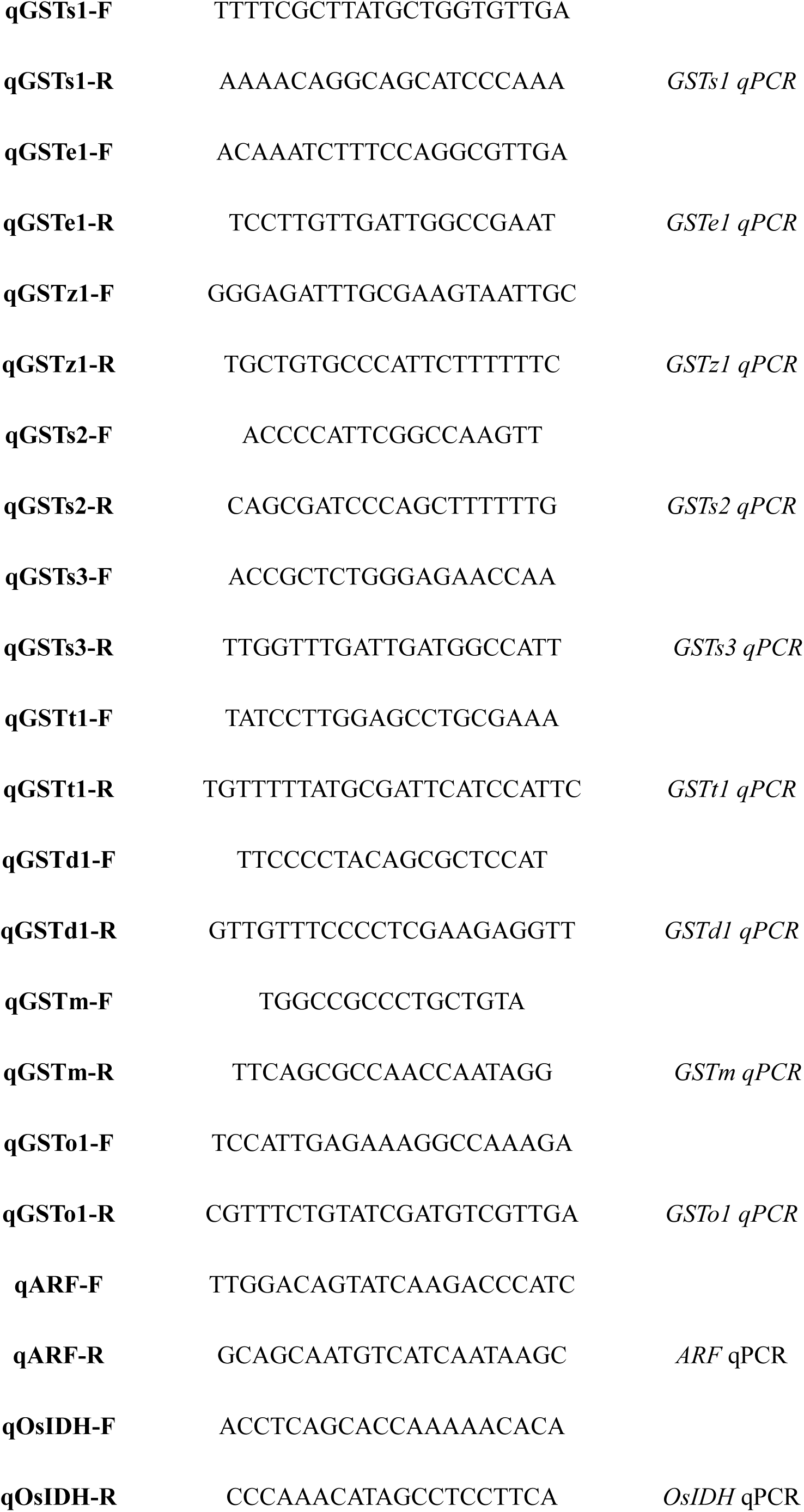
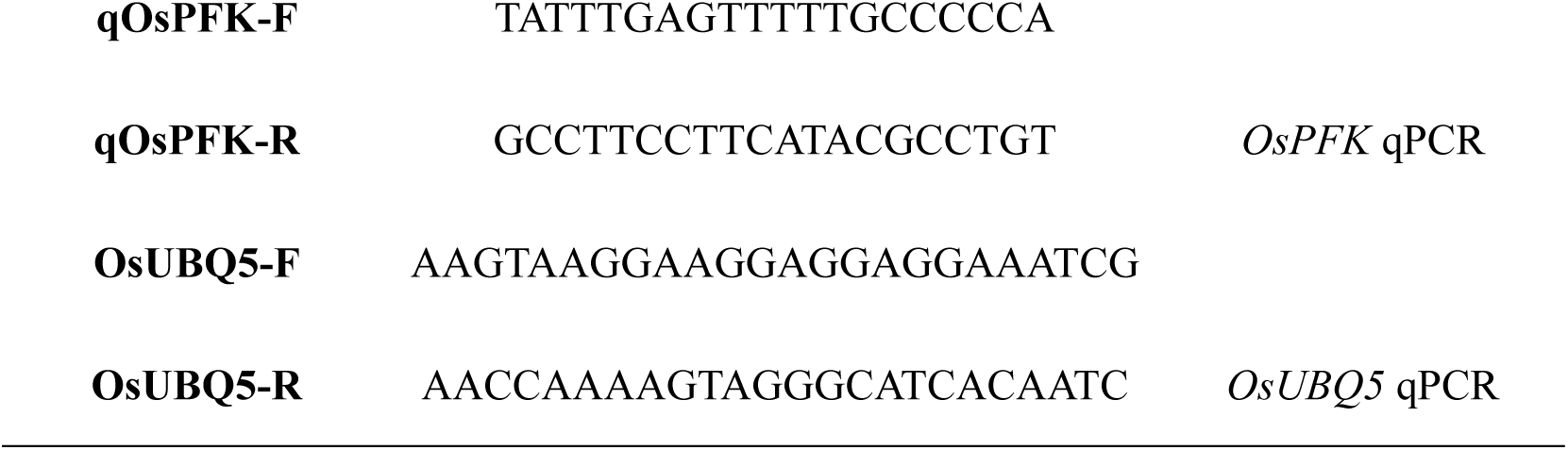
primers used for qPCR synthesis.

**Table S5.**
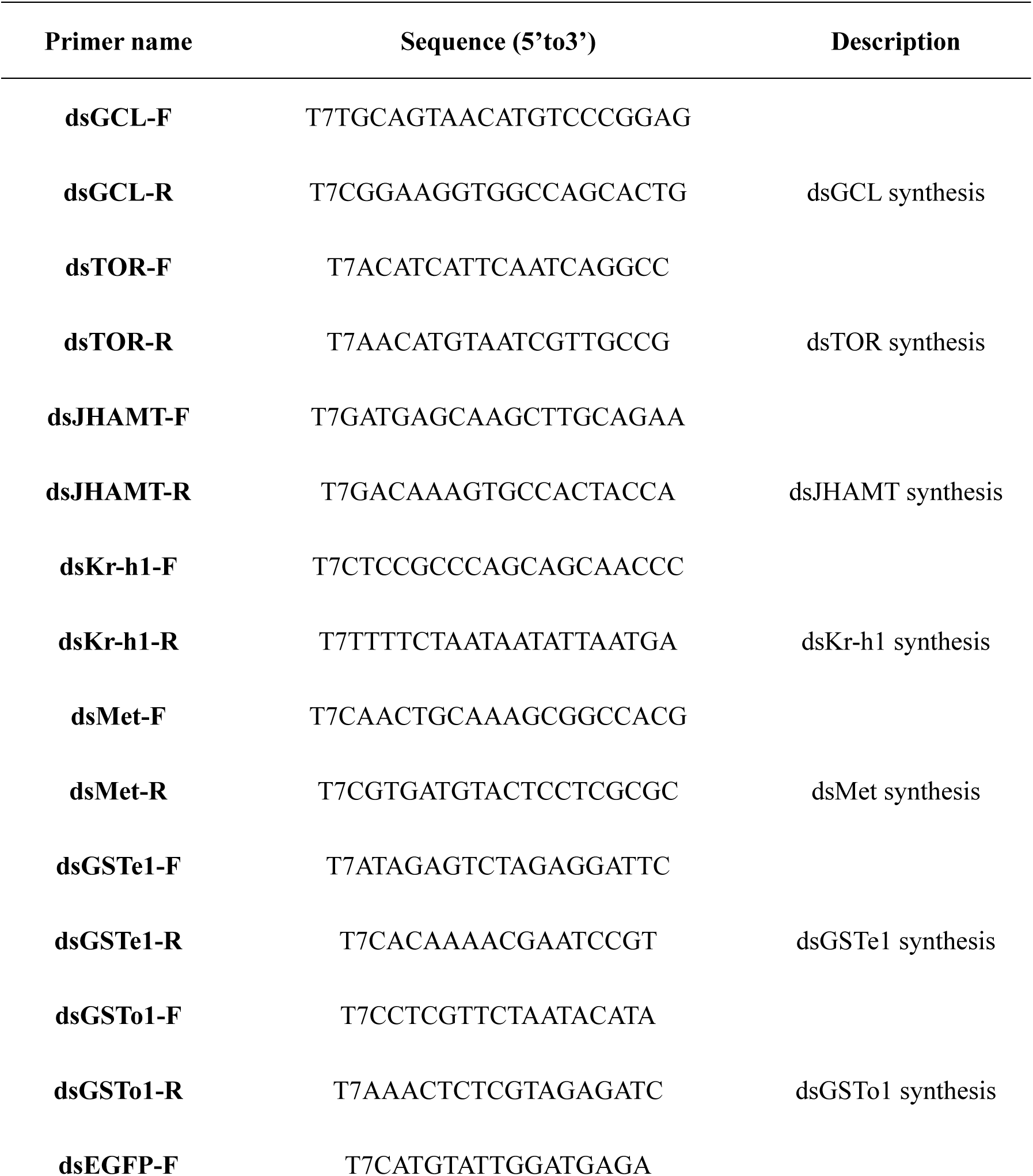
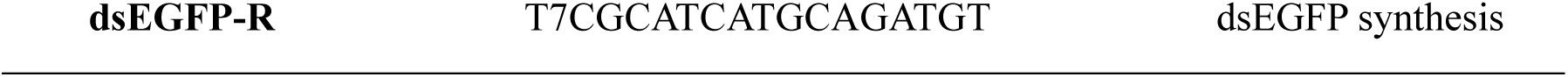
Primers used for dsRNA synthesis.

**Figure S1.**
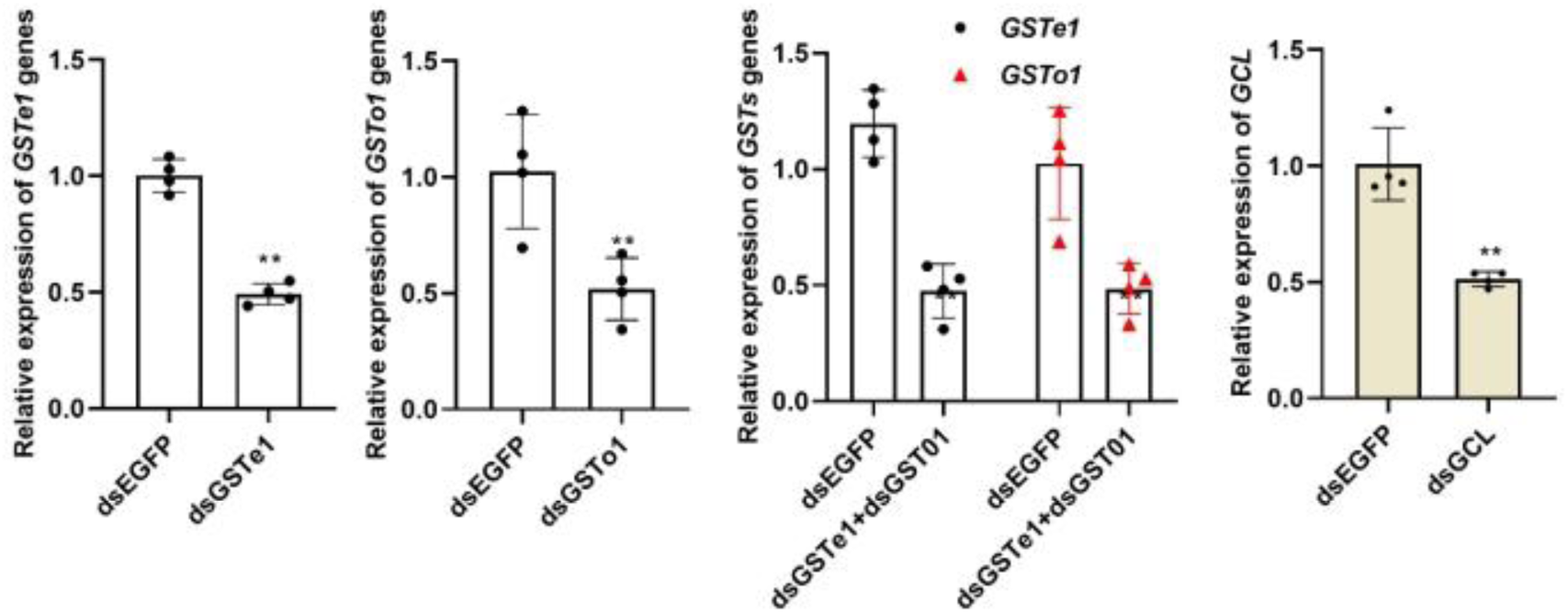
Gene expression levels five days after the third-instar nymphs were injected with the corresponding dsRNA. Data were from four biological replicate and presented as mean ± s.d.. There were analyzed by Student’s *t*-test (**\****p* < 0.05, **\*\****p* < 0.01, ns = not significant).

**Figure S2.**
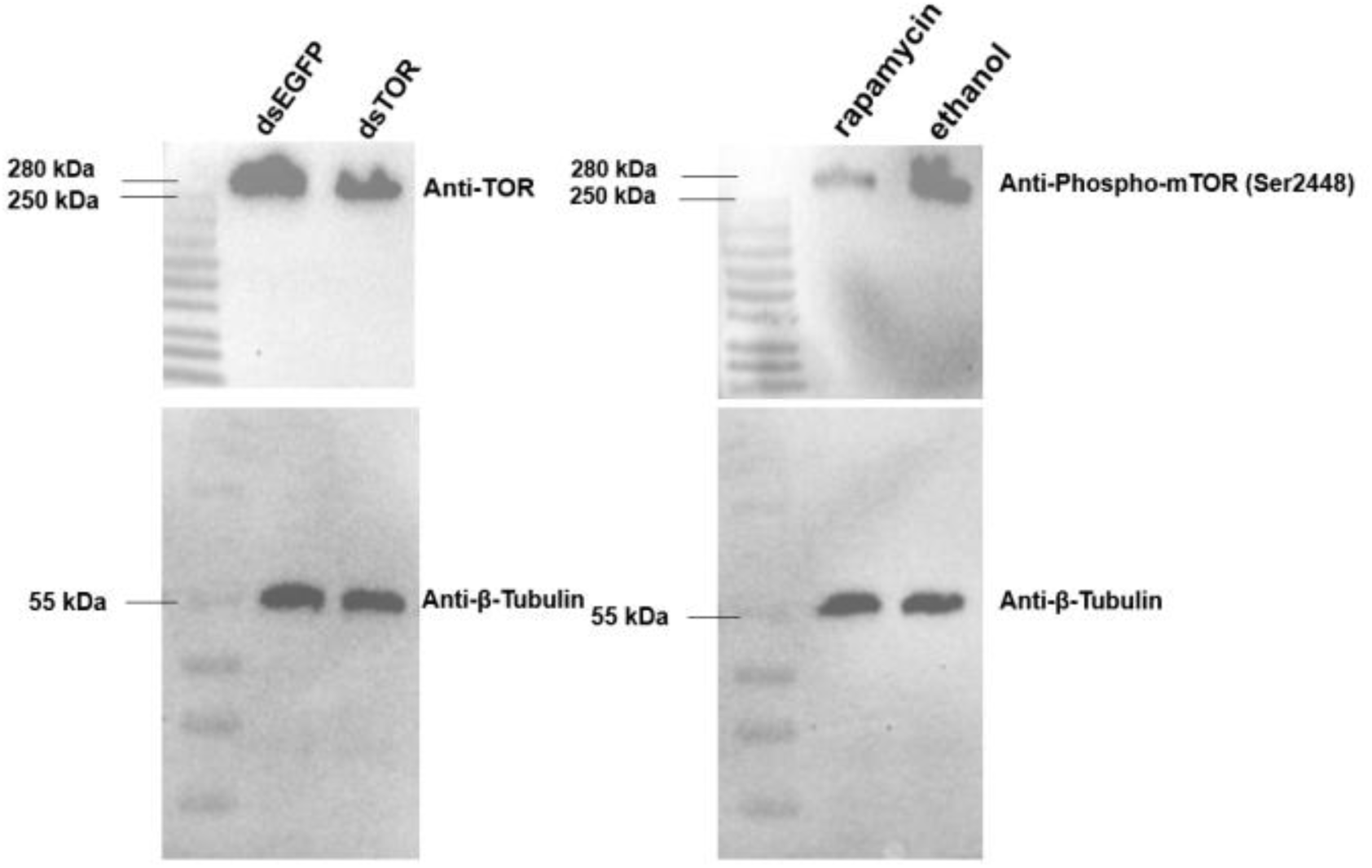
Specificity validation of Anti-TOR and Anti-Phospho-mTOR (Ser2448) antibodies. (Left) TOR protein levels were detected in third-instar nymphs at 5 days post-injection with dsTOR and reared on rice in standard IRRI solution, using an Anti-TOR rabbit polyclonal antibody (1:1000; FNab05417, FineTest, CN). Nymphs injected with dsEGFP served as the control (25 insects per group). (Right) Phosphorylation of TOR at Ser2448 was assessed in third-instar nymphs injected with rapamycin or ethanol (vehicle control) and reared under the same conditions for 5 days, using an Anti-Phospho-mTOR (Ser2448) rabbit polyclonal antibody (1:2000; AF5869, Beyotime, CN). An Anti-β-Tubulin mouse monoclonal antibody (1:2000; AF2835, Beyotime) served as a loading control. A 250 kDa Plus Prestained Protein Marker (MP202-01, Vazyme, Nanjing, CN) was used in both assays.

**Figure S3.**
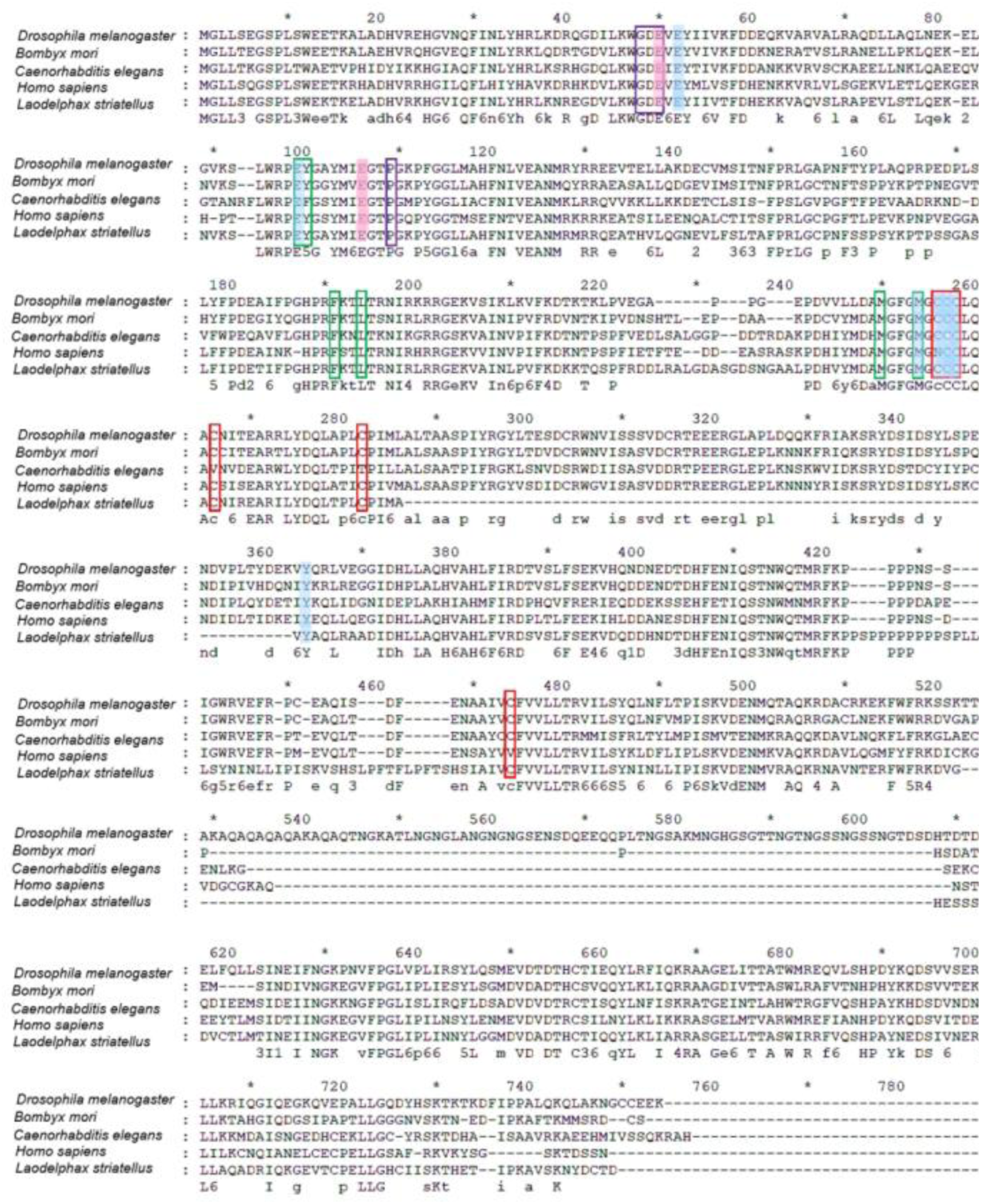
Alignment of Glutamate-cysteine ligase (GCL) amino acid sequences. The sequence of GCL in *Laodelphax striatellus* was compared to those of *Drosophila melanogaster*, *Bombyx mori*, *Caenorhabditis elegans,* and *Homo sapiens,* and the accession numbers of these proteins on NCBI are Q9W3K5.1, BDP27722.1, Q20117.2, and P48506.2 respectively. Blue shadow = glutamate-binding; Pink shadow = magnesium-binding; Purple Frame = ATP/ADP-binding; Green Frame = cysteine-binding; Red Frame = conserved cysteine

